# The Cas10 nuclease activity relieves host dormancy to facilitate spacer acquisition and retention during type III-A CRISPR immunity

**DOI:** 10.1101/2024.02.11.579731

**Authors:** Naama Aviram, Amanda K. Shilton, Nia G. Lyn, Bernardo S. Reis, Amir Brivanlou, Luciano A. Marraffini

## Abstract

A hallmark of CRISPR immunity is the acquisition of short viral DNA sequences, known as spacers, that are transcribed into guide RNAs to recognize complementary sequences. The staphylococcal type III-A CRISPR-Cas system uses guide RNAs to locate viral transcripts and start a response that displays two mechanisms of immunity. When immunity is triggered by an early-expressed phage RNA, degradation of viral ssDNA can cure the host from infection. In contrast, when the RNA guide targets a late-expressed transcript, defense requires the activity of Csm6, a non-specific RNase. Here we show that Csm6 triggers a growth arrest of the host that provides immunity at the population level which hinders viral propagation to allow the replication of non-infected cells. We demonstrate that this mechanism leads to defense against not only the target phage but also other viruses present in the population that fail to replicate in the arrested cells. On the other hand, dormancy limits the acquisition and retention of spacers that trigger it. We found that the ssDNase activity of type III-A systems is required for the re-growth of a subset of the arrested cells, presumably through the degradation of the phage DNA, ending target transcription and inactivating the immune response. Altogether, our work reveals a built-in mechanism within type III-A CRISPR-Cas systems that allows the exit from dormancy needed for the subsistence of spacers that provide broad-spectrum immunity.

## INTRODUCTION

Clustered Regularly Interspaced Short Palindromic Repeat (CRISPR) loci provide adaptive immunity to prokaryotes (Barrangou et al., 2007). These loci consist of short repetitive sequences separated by “spacer” sequences of phage or plasmid origin (Bolotin et al., 2005; Mojica et al., 2005; Pourcel et al., 2005), which are acquired from the invader’s genome upon its entry into the host (Barrangou *et al*., 2007). After being incorporated into the CRISPR locus, spacer sequences are transcribed and processed into short CRISPR RNAs (crRNAs) which associate with CRISPR-associated (Cas) proteins to mediate the base-pair recognition of nucleic acids. CRISPR-Cas systems are highly diverse and can be classified into six types (I-VI) according to their *cas* gene content (Koonin and Makarova, 2022), each of which has a different mode of immunity. Type III-A CRISPR-Cas systems are composed of a crRNA-guided complex known as Cas10-Csm (Hatoum-Aslan et al., 2013; Kazlauskiene et al., 2016) that uses the crRNA to identify complementary RNA sequences (Hale et al., 2009; Kazlauskiene *et al*., 2016). Target recognition activates two different domains of the Cas10-Csm complex: a nuclease domain (HD) that degrades ssDNA non-specifically (Kazlauskiene *et al*., 2016; Liu et al., 2019; Samai et al., 2015) and a cyclase domain (Palm) that converts ATP into cyclic tetra- or hexa-adenylates (cyclic oligoadenylates, cOA) (Kazlauskiene et al., 2017; Niewoehner et al., 2017). These are second messengers that activate CRISPR-associated Rossman fold (CARF) effectors (Stella and Marraffini, 2024). Both of these activities are transient, since Cas10 complexes are capable of cleaving the target RNA to shut off the type III response (Hale *et al*., 2009; Kazlauskiene *et al*., 2016). Type III-A CRISPR systems are commonly present in staphylococci (Cao et al., 2016; Wang et al., 2021), where they are associated with the Csm6 CARF effector, a cOA-dependent, non-sequence-specific RNase. During phage infection of *Staphylococcus aureus*, Csm6 activation is required for immunity when the target is located on a late-expressed transcript (Jiang et al., 2016) and results in the degradation of both viral and host transcripts (Rostol and Marraffini, 2019). In contrast, crRNAs that recognize early-expressed viral transcripts do not require this RNase and presumably rely on the ssDNase activity of the Cas10-Csm complex to provide immunity. In the case of spacers that target plasmid transcripts, RNA cleavage by Csm6, as well as its ortholog Csx1 (present within type III-B CRISPR loci in other organisms), results in a growth arrest of the cell (Rostol and Marraffini, 2019; Sheppard et al., 2016).

Whether Csm6 activation during the type III-A CRISPR response against phage infection results in the growth arrest of host and, if so, whether there are mechanisms that enable exit from dormancy is not known. Perhaps more importantly, how the growth arrest affects spacer acquisition and maintenance is a central question in CRISPR immunity, as not only type III, but also type VI (Abudayyeh et al., 2017; Meeske et al., 2019), CRISPR systems are capable of mediating cell dormancy. Hosts that acquire spacer sequences capable of mediating a halt to their replication would be at a disadvantage with uninfected cells that continue growing or with other bacteria that acquire spacers that protect without triggering cell dormancy. Moreover, even if dormancy-inducing spacers are acquired and fixed in the population, how they subsist after an infection event that triggers the growth arrest of the host they reside in is not known. Spacer acquisition into the *Thermus thermophilus* type III-A CRISPR-Cas system after phage infection has been studied previously in detail (Artamonova et al., 2020), however whether immunity triggers dormancy and/or whether the CARF effectors present in the strain used (HB27c, which harbors three CARF genes) are required for immunity, was not investigated. Here we thoroughly characterized the Csm6 response to phage infection in staphylococci. We found that Csm6 is not only required for defense when the target transcript is expressed late in the phage lytic cycle, but also when it is mediated by crRNAs that anneal to early-expressed viral RNAs. Targeting of late transcripts generated a prolonged dormancy of the infected cells that prevented the propagation not only of the target phage but also of other viruses present in the environment that fail to replicate in the arrested cells. We found that when immunity requires Csm6 activity, the ssDNase activity of the Cas10-Csm complex is required to enable the regrowth of the infected cells. Our results reveal a built-in mechanism within type III-A CRISPR-Cas systems that allows the exit from dormancy needed for the retention of spacers that provide broad-spectrum immunity.

## RESULTS

### Type III-A CRISPR spacers targeting late-expressed ΦNM4γ4 transcripts induce growth arrest of the infected cell

While the induction of host dormancy after activation of Csm6 has been studied when plasmids trigger the type III-A CRISPR-Cas defense (Rostol and Marraffini, 2019), little is known about how Csm6-mediated dormancy affects immunity against phages. Previous work investigated the requirement of Csm6 for efficient defense using only a limited number of select spacers that targeted the staphylococcal phage ΦNM1γ6 (Goldberg et al., 2014; Jiang *et al*., 2016). Therefore, we decided to test the immunity mediated by a library of spacers that covered the complete sequence of the ΦNM4γ4 phage (Bae et al., 2006; Heler et al., 2015) (Fig. S1A), a close relative of ΦNM1γ6 (Bae *et al*., 2006; Banh et al., 2023) which we have successfully used for high throughput CRISPR targeting studies (Kenney and Marraffini, 2023). To construct this library, we designed 40,338 spacers that matched both strands of the genome every two nucleotides. Oligonucleotides harboring the different spacer sequences were amplified and cloned into a type III-A CRISPR array present within the shuttle vector pLZ12 (Perez-Casal et al., 1991). The plasmid spacer library was introduced into *Staphylococcus aureus* RN4220 (Kreiswirth et al., 1983) cells harboring a spacer-less *S. epidermidis* type III-A CRISPR-Cas system (Fig. S1B) cloned into the staphylococcal vector pC194 (Horinouchi and Weisblum, 1982) (pNA1, Table S6). Library plasmids were isolated from cultures and spacers were amplified and analyzed by next generation sequencing (NGS) to determine their abundance. We were able to detect 40,037 unique spacer sequences, with equivalent frequency of reads for the great majority of them (Fig. S1C, the expected frequency is 1/40,338 or 2.4 × 10^-5^). We then infected the cultures with ΦNM4γ4, used NGS to measure the abundance of each spacer 5 and 24 hours after phage addition, and calculated enrichment ratios as the read count post-infection divided by its abundance in the uninfected library (Figs. 1A and S1D). As expected from the RNA targeting of the Cas10 complex (Elmore et al., 2016; Estrella et al., 2016; Kazlauskiene *et al*., 2017; Kazlauskiene *et al*., 2016; Niewoehner *et al*., 2017; Samai *et al*., 2015), the data for both timepoints showed little to no enrichment for spacers matching the plus strand sequences of ΦNM4γ4, which, based on the direction of the open reading frames, should not be transcribed (Fig. S1A). On the other hand, for spacers generating a crRNA complementary to predicted transcripts, matching the minus strand of the phage genome, we found that only those targeting a region spanning ∼1 to 15 kb of ΦNM4γ4 were highly enriched at both timepoints post-infection, with a decrease in spacer accumulation from 15 to 18 kb (Figs. 1A and S1D). This is a somewhat surprising result given that the rest of the genome (18 to 42 kb) must be transcribed during the lytic cycle to generate RNA targets for the Cas10-Csm complex, and therefore should also be enriched. Indeed, a previous study showed that targeting of ΦNM4γ4 using the *S. epidermidis* type III-A CRISPR system programmed with a crRNA complementary to the *gp47* transcript (located at approximately the 20 kb position in the genome), which encodes the major head protein and therefore it is expected to be expressed late in the lytic cycle, reduced plaque formation to undetectable levels (Pyenson et al., 2017). To clarify this, we performed RNA-seq analysis at 5, 15 and 30 minutes after ΦNM4γ4 infection (Fig. S1E and Table S2). We found that transcription of the phage genome occurs primarily from two promoters; an early promoter (PE, Fig. S1A) that directs the expression of the 1-15 kb region during the first five minutes after infection, and a second late promoter (PL) responsible for the transcription of the 15-42 kb region, starting sometime between 5 and 15 minutes after infection. These results indicate that the spacers most enriched from the library are those that target early-expressed ΦNM4γ4 transcripts. Indeed, we found a strong correlation between spacer enrichment ratios and target RNA-seq reads obtained 5 minutes post-infection (Fig. S1F).

**Figure 1.**
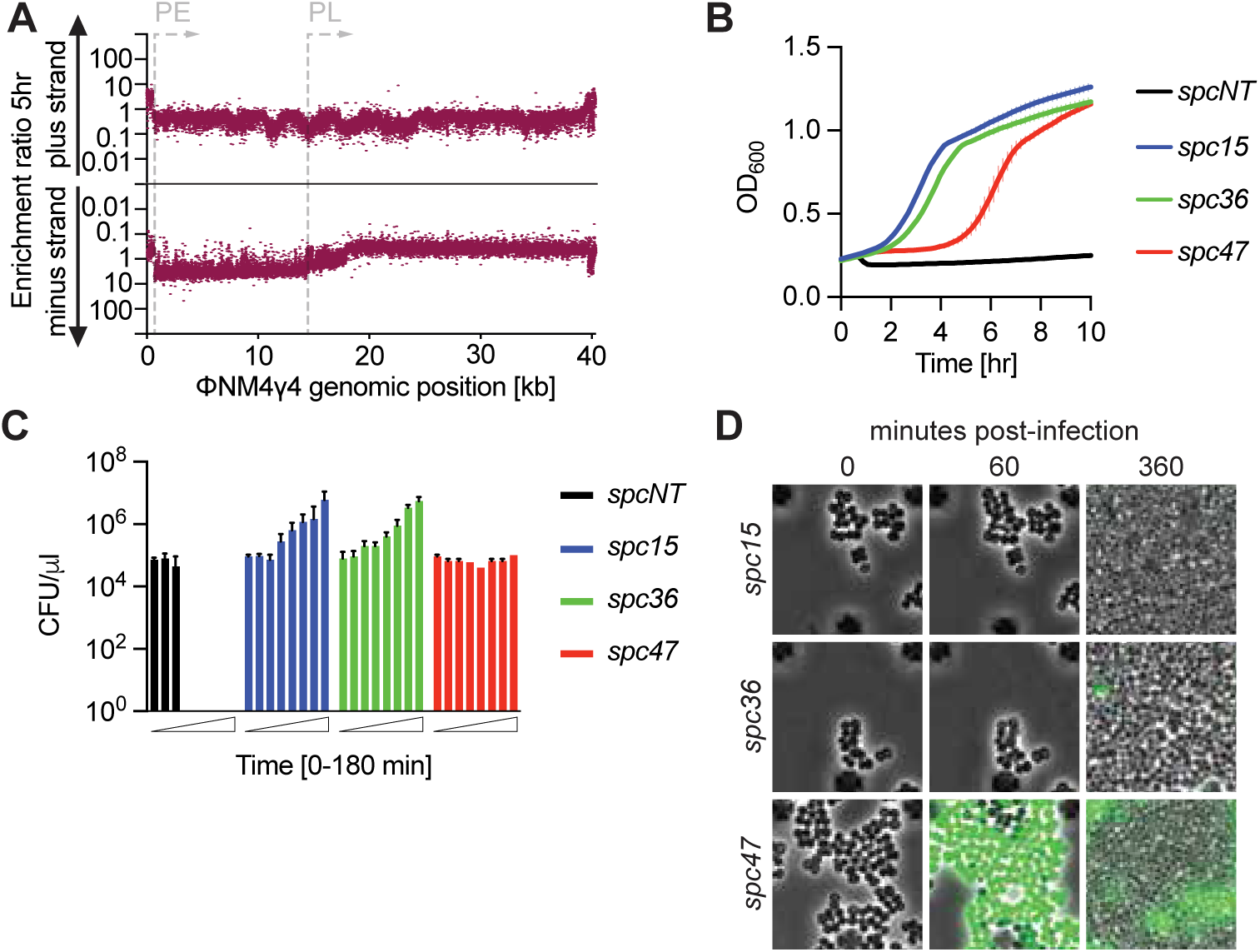
Type III-A CRISPR immunity induces the growth arrest of the infected cell. **(A)** Enrichment ratio of spacers targeting the plus or minus strands of the ΦNM4γ4 DNA 5 hours after phage infection of staphylococci carrying pCRISPR, plotted according to their genomic position. **(B)** Mean (± SD, n = 3 biological replicates) OD_600_ values of staphylococcal cultures harboring a pCRISPR plasmid programmed with the indicated spacers (*spc15, spc36, spc47,* Δ*spc*) after infection with ΦNM4γ4 at MOI 10. **(C)** Mean (± SD, n = 3 biological replicates) CFU/µl present in cultures harboring pCRISPR programmed with the indicated spacers (*spc15, spc36, spc47,* Δ*spc*), with samples taken every 30 minutes for three hours, after infection with ΦNM4γ4 at an MOI of 10. **(D)** Time-course fluorescence microscopy of staphylococci carrying pCRISPR plasmids programmed with different spacers, 0, 60 and 360 minutes after infection with ΦNM4γ4^GFP^.

Our previous work using select spacers against ΦNM1γ6 demonstrated that Csm6 is required for immunity mediated by spacers that target late-, but not early-expressed viral genes (Goldberg *et al*., 2014; Jiang *et al*., 2016). In addition, we also found that when type III-A CRISPR immunity is triggered by a plasmid expressing a target transcript, the activation of Csm6 leads to a growth arrest of the host cell (Rostol and Marraffini, 2019). Therefore, we hypothesized that the library spacers matching the 18-42 kb region trigger the activation of this nuclease and therefore cause a growth arrest of staphylococci harboring them that prevents their enrichment. To test this, we cloned spacer sequences matching PE and PL transcripts into pCRISPR. Plasmids were introduced into *S. aureus* RN4220 cells and growth of the cultures was evaluated by optical density at 600 nm (OD_600_). After treatment with ΦNM4γ4 at a multiplicity of infection (MOI) of 10, control cells carrying a CRISPR system without a spacer (Δ*spc*), lysed rapidly (Fig. 1B). In contrast, spacers targeting PE-derived transcripts (*spc15* and *spc36*) enabled the growth of the infected cultures (Fig. 1B). The strain harboring the spacer targeting a transcript expressed from the PL promoter (*spc47*), however, also survived phage infection but displayed a growth defect (Fig. 1B). To corroborate this growth arrest, we enumerated colony forming units (CFU) within the four cultures at different times after infection with ΦNM4γ4 at MOI 10 (Fig. 1C). Due to phage lysis, CFU counts drastically decreased in the absence of a targeting spacer. For cultures harboring *spc15* and *spc36*, which did not display a growth delay after infection, CFU values steadily increased. However, CFUs for the *spc47*-harboring staphylococci remained constant, at least for the first 3 hours after infection, a result that confirms the arrest of the infected cells carrying this spacer.

In spite of this cessation in proliferation, *spc47* cultures grow to high densities after infection (Fig. 1B). To investigate this in more detail, we used a virus that expresses *green fluorescent protein* (ΦNM4γ4*^gfp^*, Fig. S1A) and followed bacterial proliferation using fluorescence microscopy. We found that while staphylococci lacking a targeting spacer (Δ*spc*) displayed green fluorescence and lysed (Fig. S1G and Supplementary b s 1-2), cultures harboring *spc15* continued their growth normally after infection, without detection of green fluorescence nor lysis (Fig. 1D). Similarly, very few *spc36* cells turned green after infection, emitting only dim fluorescence, with most staphylococci displaying a minor division stall after infection that was followed by vigorous proliferation (Fig. 1D). In contrast, cells producing the *spc47* crRNA turned green and stopped dividing, with non-infected cells (non-green, Fig. 1D) restarting growth at about 90 minutes post-infection to take over the bacterial population. We conclude that spacers targeting late-expressed phage transcripts mediate type III-A CRISPR immunity through the generation of a growth arrest that prevents viral propagation to promote the survival of the uninfected cells of the culture. Therefore, although dormancy-promoting spacers are important to provide defense, they impose a fitness cost for the cells that harbors them that results in their depletion from the infected population. In contrast, spacers that target PE-derived transcripts enable the replication of the infected staphylococci, conferring a fitness advantage during phage infection that leads to their accumulation in the bacterial culture.

### Immunity triggered by early-expressed transcripts also requires Csm6 but does not induce a growth arrest

The lack of growth arrest after infection of cells that carry spacers targeting PE-derived transcripts, as well as their positive selection in our library experiments, suggest that these spacers do not trigger Csm6. However, our current understanding of the molecular events that occur during type III-A CRISPR immunity indicate that, since recognition of a complementary RNA sequence results in the synthesis of cOA molecules by the Cas10 complex (Kazlauskiene *et al*., 2017; Niewoehner *et al*., 2017), spacers targeting PE-derived transcripts should also induce the production of second messengers that activate Csm6. Indeed, we did observe a minor growth defect during infection for staphylococci harboring *spc36*, detected both by optical density measurements (Fig. 1B) as well as microscopy experiments. To study a possible role for Csm6 activation during type III-A targeting of early-expressed ΦNM4γ4 transcripts, we performed infection experiments in the absence of this RNase, introducing the spacer library into staphylococci carrying a plasmid-borne type III-A system that expresses a catalytically dead Csm6, dCsm6 (Jiang *et al*., 2016; Niewoehner *et al*., 2017; Niewoehner and Jinek, 2016) (pNA23, Fig. S1B and Table S6). We were able to detect 39,953 spacer sequences after NGS of the resulting culture (Fig. S2A). As in the previous experiment, spacers producing crRNAs not complementary to the RNA generated during ΦNM4γ4 infection (matching the plus strand) or targeting PL-derived transcripts were not enriched due to their inability to confer immunity or to their requirement for Csm6 to provide defense, respectively; both at 5 and 24 hours after phage treatment (Figs. 2A and S2B). Surprisingly, approximately half of the spacers targeting early-expressed transcripts also displayed lack of selection (Fig. 2A). This result suggested that these spacers require the activation of Csm6 to confer immunity. Indeed, the number of unique spacer sequences matching the minus strand that we were able to detect 24 hours after phage selection decreased to 9,186 in the presence of dCsm6 from 19,859 in cultures carrying a wild-type III-A CRISPR system, with most of the missing spacers mapping to the region with low enrichment (Figs. S2C-D). This depletion suggests that many of the infected cells harboring these spacers undergo lysis in the absence of Csm6 activity. To corroborate this, we followed the growth of cultures carrying a single spacer, cloned into a version of pCRISPR that expresses dCsm6, infected with ΦNM4γ4 at a MOI of 10 (Fig. 2B). We found that *spc15* mediated complete defense, equivalent to that observed in the presence of wild-type Csm6 (Fig. S2E). In contrast, staphylococci carrying *spc47* succumbed to phage infection, similarly to cells lacking a targeting spacer, a result in accordance with the importance of the Csm6-induced growth arrest for the survival of the population. These observations were further corroborated both by CFU enumeration after infection (Fig. 2C) and fluorescent microscopy of ΦNM4γ4*^gfp^*-infected cells (Fig. 2D), with both experiments demonstrating the death of the *spc47*-d*csm6* staphylococci.

**Figure 2.**
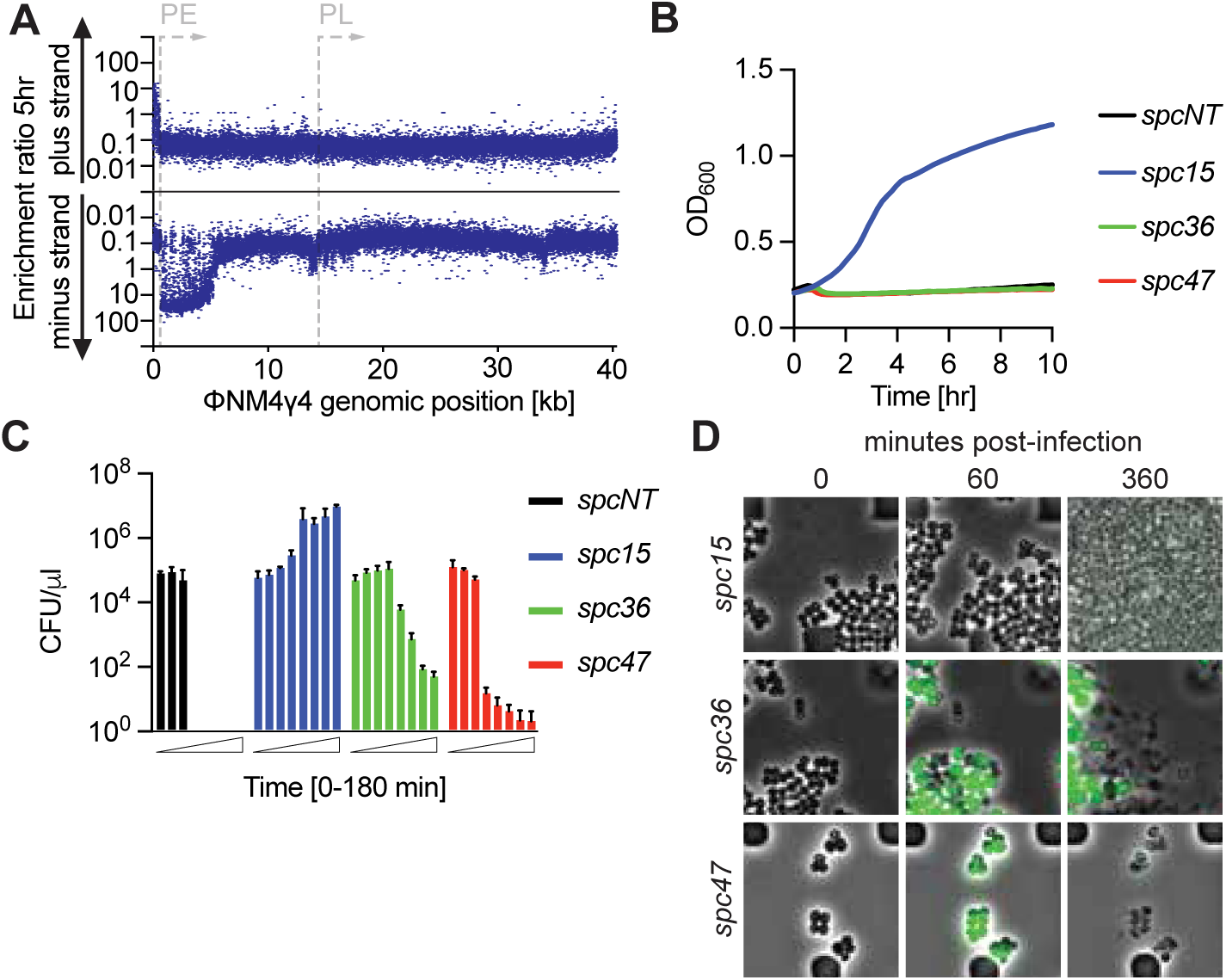
Spacers can require Csm6 for immunity without inducing a growth arrest. **(A)** Enrichment ratio of spacers targeting the plus or minus strands of the ΦNM4γ4 DNA 5 hours after phage infection of staphylococci carrying pCRISPR(d*csm6*), plotted according to their genomic position. **(B)** Mean (± SD, n = 3 biological replicates) OD_600_ values of staphylococcal cultures harboring a pCRISPR(d*csm6*) plasmid programmed with the indicated spacers (*spc15, spc36, spc47,* Δ*spc*) after infection with ΦNM4γ4 at MOI 10. **(C)** Mean (± SD, n = 3 biological replicates) CFU/µl present in cultures harboring pCRISPR(d*csm6*) programmed with the indicated spacers (*spc15, spc36, spc47,* Δ*spc*), with samples taken every 30 minutes for three hours, after infection with ΦNM4γ4 at an MOI of 10. **(D)** Time-course fluorescence microscopy of staphylococci carrying pCRISPR(d*csm6*) plasmids programmed with different spacers, 0, 60 and 360 minutes after infection with ΦNM4γ4^GFP^.

Consistent with the lack of strong selection after infection of the dCsm6 library of spacers targeting the downstream region of the PE-derived ΦNM4γ4 transcript (Fig. 2A), *spc36* failed to provide immunity in the absence of Csm6 activity, as measured by optical density (Fig. 2B), CFU enumeration (Fig. 2C) and fluorescence microscopy (Fig. 2D), the latter showing the lysis of the infected cells. Interestingly, the decrease in CFU counts is much less drastic than in Δ*spc* control cells, a result that indicates that either the sequence-specific target transcript binding, target RNA cleavage, and/or non-specific DNA degradation of the Cas10 complex can interfere with the phage lytic cycle. In addition, we tested other spacer sequences in this area (*spc29* and *spc35*, Fig. S1A) and found similar growth curves to that of *spc36* (Fig. S2F). Altogether these results demonstrate that spacers that target the downstream region of the early-expressed transcript of ΦNM4γ4 (“PE-downstream” spacers) require Csm6 activity to provide defense. However, as opposed to the Csm6-dependent response of spacers targeting the late-expressed transcript (*spc47*), PE-downstream spacers do not trigger a growth arrest, suggesting the existence of a mechanism to enable the proliferation of staphylococci in which Csm6 has been activated, and therefore their enrichment within the infected bacterial population.

### Cas10 ssDNase activity is required to prevent Csm6-dependent dormancy when the target is located on an early-expressed transcript

The lack of GFP detection in the majority of *spc15*- and *spc36*-harboring staphylococci after infection suggests that the CRISPR response within these cells can prevent the transcription of the ΦNM4γ4 genome, even of sequences expressed early in the lytic cycle. Previously we showed that the nuclease activity of Cas10 is required to eliminate a target plasmid DNA during *S. epidermidis* type III-A CRISPR-Cas immunity (Rostol and Marraffini, 2019). Therefore, we hypothesized that this activity could be important for inactivating the viral DNA in infected cells equipped with spacers targeting early-expressed ΦNM4γ4 transcripts. To test this, we performed infection experiments in staphylococci carrying a mutation in *cas10* that abrogates nuclease activity, Cas10^HD^ (Kazlauskiene *et al*., 2016; Rostol and Marraffini, 2019) (pNA24, Fig. S1B and Table S6). We then introduced the spacer library into these cells, being able to detect 40,079 spacer sequences after evaluation of the resulting culture by NGS (Fig. S3A). As before, spacers producing crRNAs not complementary to the RNA generated during ΦNM4γ4 infection (matching the plus strand) were not enriched, neither at 5 (Fig. 3A) nor at 24 hours (Fig. S3B) after phage treatment. This was also the case for spacers producing functional crRNAs, with slightly more abundance for the those mediating the targeting of PL-derived transcripts (Fig. 3A; average enrichment of spacers targeting PE-transcripts, 0.78±0.26; of PL-derived transcripts, 1.96±0.73). We hypothesize that this is a consequence of the rapid activation of Csm6 by crRNAs complementary to PE-transcripts, which leads to an earlier growth arrest (and therefore less cell proliferation) than those that target PL-derived phage RNAs.

**Figure 3.**
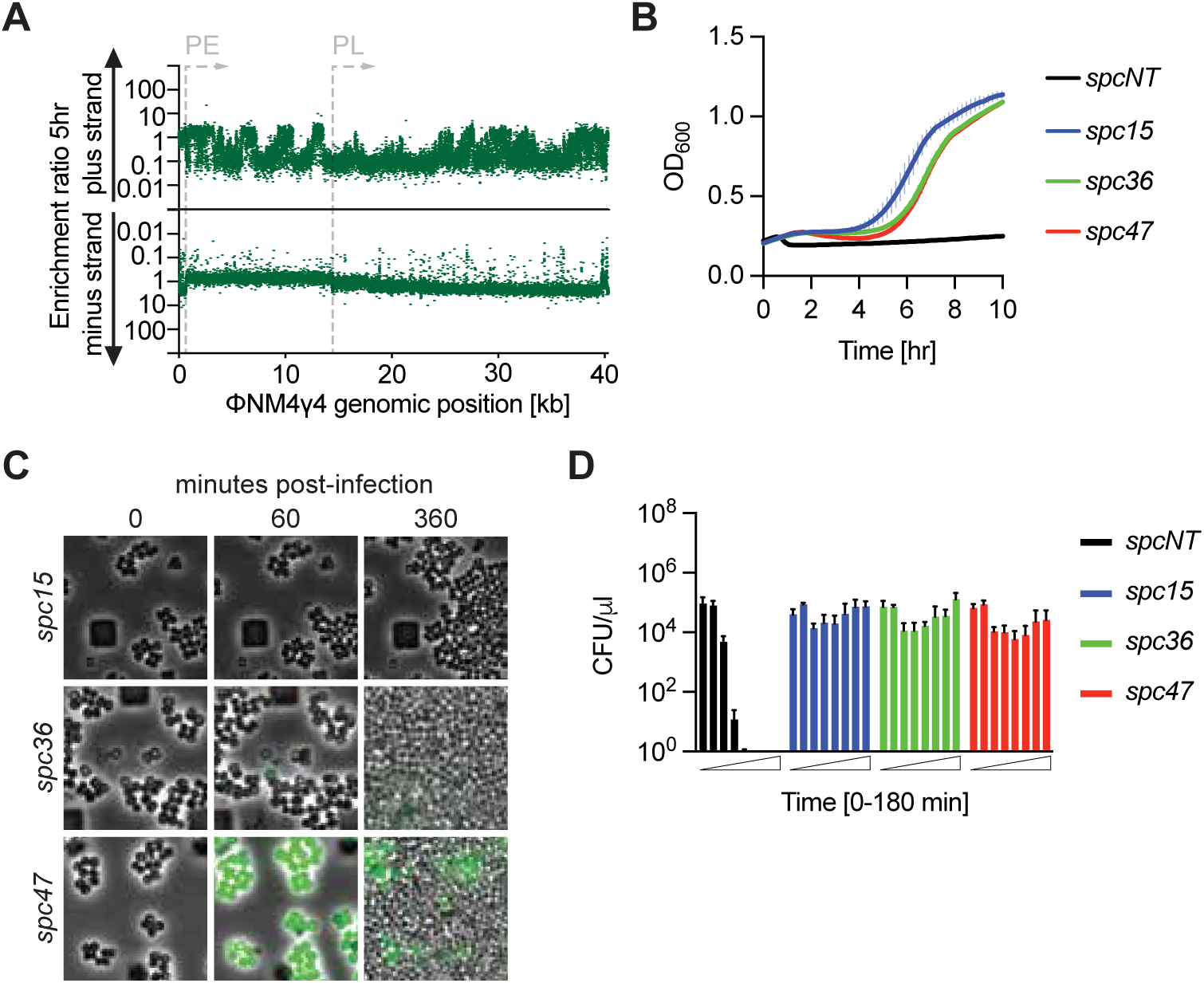
Cas10 ssDNase activity can relieve Csm6-dependent dormancy. **(A)** Enrichment ratio of spacers targeting the plus or minus strands of the ΦNM4γ4 DNA 5 hours after phage infection of staphylococci carrying pCRISPR(*cas10*^HD^), plotted according to their genomic position. **(B)** Mean (± SD, n = 3 biological replicates) OD_600_ values of staphylococcal cultures harboring a pCRISPR(*cas10*^HD^) plasmid programmed with the indicated spacers (*spc15, spc36, spc47,* Δ*spc*) after infection with ΦNM4γ4 at MOI 10. **(C)** Time-course fluorescence microscopy of staphylococci carrying pCRISPR(*cas10*^HD^) plasmids programmed with different spacers, 0, 60 and 360 minutes after infection with ΦNM4γ4^GFP^. **(D)** Mean (± SD, n = 3 biological replicates) CFU/µl present in cultures harboring pCRISPR(*cas10*^HD^) programmed with the indicated spacers (*spc15, spc36, spc47,* Δ*spc*), with samples taken every 30 minutes for three hours, after infection with ΦNM4γ4 at an MOI of 10.

Given that Csm6 should be activated in the absence of Cas10 nuclease activity, all spacers should be able to provide the population-level immunity mediated by this RNase. To test this, we performed phage infection experiments of mono-cultures harboring a single spacer and the *cas10*^HD^ allele at MOI 10. In the presence of *spc15*, *spc36* or *spc47*, addition of ΦNM4γ4 resulted in a delay in the increase of OD_600_ (Fig. 3B), remarkably similar to the growth curve of cultures harboring *spc47* in a wild-type *cas10* background (Fig. S3C), for which immunity depends on Csm6. Indeed, the same experiment in the presence of *cas10*^HD^/d*csm6* double mutant showed the complete absence of defense, with cultures collapsing upon phage addition (data not shown). To investigate in more detail the growth delay of the *cas10*^HD^ cultures, we followed cells under the microscope after the addition of ΦNM4γ4*^gfp^*. We observed that staphylococci harboring *spc15* did not express GFP after infection, however they stalled their proliferation after infection (Fig. 3C). Similarly, cells equipped with *spc36* showed very dim and non-uniform green fluorescence and also stopped growing after phage infection (Fig. 3C). We hypothesize that the lack of green fluorescence signal is a consequence of early Csm6-mediated degradation of the *gfp* phage transcript, an interpretation that is in line with the lower number of spacer reads for the spacers that target PE-derived spacers (Fig. 3A), for which Csm6 activation stops the growth of the cells that harbors them shortly after infection. In contrast, staphylococci that harbored *spc47* in the absence of Cas10 nuclease activity showed strong green fluorescence during their growth arrest (Fig. 3C), similarly to the cells that carry wild-type *cas10* (Fig. 1D), with uninfected cells that do not express GFP taking over the bacterial population. Finally, the growth arrest detected in both growth curves and microscopy was corroborated by enumerating CFUs at different time-points after infection at MOI 10 an experiment that showed a constant number of cells over time, similar for all three spacer sequences (Fig. 3D). Altogether these experiments confirmed that the immunity mediated by *spc47* has identical properties in the presence or absence of the Cas10 nuclease activity, a result that demonstrates that this immunity is mediated exclusively by Csm6. In contrast, the Cas10 nuclease is fundamental for the immunity provided by *spc15* and *spc36*, and promotes the proliferation of the infected cells. When this activity is eliminated, these spacers trigger Csm6 and convey defense at the population level, triggering the arrest of the infected cell to prevent phage proliferation. In the case of the immunity provided by *spc36*, and possible other spacers that mediate targeting of the downstream region of the PE transcript, the prolonged dormancy observed in the presence of *cas10*^HD^ demonstrates that Cas10 ssDNase activity is necessary for allowing the regrowth of the infected cells carrying these spacers.

### Spacer acquisition during the type III-A CRISPR response is biased against sequences that mediate growth arrest

Our experiments indicate that spacers that target PE-derived transcripts provide a selective advantage to the cell that contains them, owing to the ssDNase activity of Cas10 that allows the normal growth of the infected host, presumably through a direct attack of the viral genome. In contrast, spacers that generate crRNAs complementary to PL-transcribed phage RNA molecules are at a disadvantage, since they trigger the arrest of the host that harbors them and therefore constitute spacers that do not accumulate in the bacterial population. In principle, these results should extend to the process of spacer acquisition during the type III-A CRISPR-Cas response against phages, i.e., lead to a robust acquisition of PE-targeting spacers and a very low frequency of acquisition for PL-targeting ones. To test this, we infected cells harboring a pCRISPR plasmid with a single repeat and no spacers (Aviram et al., 2022) with ΦNM4γ4 at an MOI of 2 in semi-solid agar plates, conditions that were previously shown to minimize the rise of non-CRISPR, phage-resistant colonies (Pyenson and Marraffini, 2020). The surviving colonies were collected to extract plasmid DNA and subject it to NGS to identify the newly acquired spacers. Mapping of the spacer sequences to the phage genome revealed that the vast majority were inserted in the orientation that, when the CRISPR array is transcribed, would generate a crRNA complementary to PE-generated phage RNA (Fig. 4A, “minus strand” spacers, and Table S1). This pattern is similar to that obtained with our spacer library (Fig. 1A), where only the sequences that confer immunity to the individual infected cells but not those that mediate a growth arrest, are enriched. Therefore, this result suggests that acquired spacer sequences are selected for their ability to provide immediate defense.

**Figure 4.**
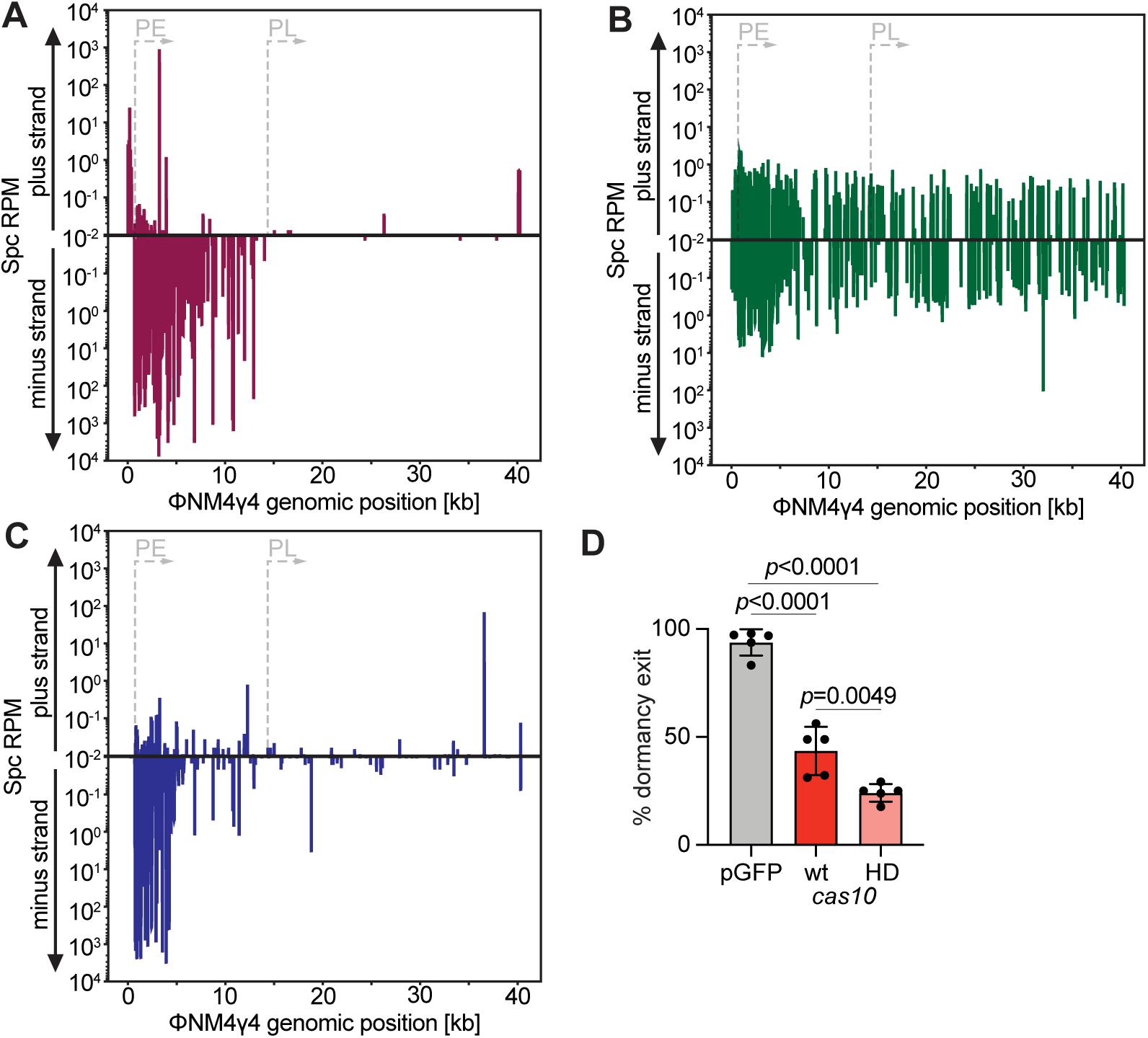
Spacers that mediate growth arrest are rarely acquired. **(A)** Abundance (in reads per million, RPM) of spacer sequences acquired by staphylococci harboring pCRISPR after infection with ΦNM4γ4. Spacers matching the plus or minus strands of the viral DNA are plotted according to their position in the phage genome. **(B)** Same as **(A)** but following infection of staphylococci harboring pCRISPR(*cas10*^HD^). **(C)** Same as **(C)** but following infection of staphylococci harboring pCRISPR(d*csm6*). **(D)** Mean (± SD, n=5 independent experiments) percentage of single cells infected with ΦNM4γ4*^gfp^*that grew after index-sorting into a 96-well plate. Uninfected staphylococci expressing GFP on a plasmid (pGFP) were used as control. The growth of green-fluorescent, single cells obtained from staphylococcal cultures harboring pCRISPR or pCRISPR(*cas10*^HD^) programmed with *spc47* was compared. *p* values were calculated with One-Way ANOVA and Tukey’s multiple comparisons tests.

Alternatively, there could be a promoter-specific effect on spacer acquisition, where sequences from the DNA transcribed from PE, but not those from the DNA transcribed by PL, are uniquely captured by the Cas1/2 integrase complex (Nunez et al., 2015; Yosef et al., 2012). Transcription-dependent spacer acquisition was found in both *S. aureus* and *Streptococcus thermophilus* that harbor type III-A systems, and leads to the preferential incorporation of spacers from rDNA and tDNA genes (Aviram *et al*., 2022; Zhang et al., 2021). To explore these different possibilities, we performed spacer acquisition experiments using pCRISPR carrying the *cas10*^HD^ allele, a genetic background in which spacers that produce crRNAs that target both PE- and PL-derived viral transcripts mediate a growth arrest and population-level immunity (Figs. 3B-D). We found that spacer acquisition occurred at a very low frequency, from both the minus and plus strand, and therefore most likely the data collected represents background NGS reads (Fig. 4B and Table S1). This data demonstrates that there is a severe bias against the incorporation spacers that mediate growth arrest, regardless of how and/or when their targets are transcribed. Finally, to corroborate the importance of the Cas10-mediated immunity in spacer acquisition, we performed experiments in the d*csm6* genetic background, where only spacers matching the upstream region of the PE operon provide strong defense in our library analysis (Fig. 2A). The distribution of acquired spacers was similar to that of the enrichment within the spacer library after infection, i.e., the majority of spacers were acquired form the upstream region of the PE-transcribed operon (minus strand, Fig. 4C, and Table S3). Altogether these results demonstrate that spacer acquisition by type III-A CRISPR systems displays a strong preference for sequences that mediate immunity through Cas10’s ssDNase activity and enable the proliferation of the infected cells.

### Cas10’s ssDNase activity is required to relieve the dormancy of infected cells

Interestingly, a very low number (39) of spacers were acquired from the PL-transcribed region of the phage (Table S3) during the wild-type type III-A CRISPR-Cas response, even if they are not able to activate the ssDNase activity that is required for the successful capture of new spacers that we described above. In addition, previous bioinformatic analysis that described type III-A CRISPR loci in environmental and clinical staphylococcal isolates (Cao *et al*., 2016; Wang *et al*., 2021) reported the presence of spacers that are predicted to target late-expressed genes (Table S4). In order to maintain the dormancy-inducing spacers in the population, during and after their acquisition, non-proliferating cells should be able to resume growth sometime after infection. We looked for this by following cells with green fluorescence, to determine if they can restart their replication. However, the vigorous regrowth of the uninfected cells prevented us from distinguishing the fate of individual cells. Therefore, we subjected *spc47* cultures (Fig. 1D) to fluorescently activated single-cell sorting into 96-well plates to monitor the growth of individual infected cells. Uninfected cultures of mixed cells carrying a GFP- or mCherry-expressing plasmid were used as a control to ensure that the process of single-cell sorting did not negatively impact cell viability and to validate that a single cell is sorted into each well (Figure S4A-B). Surprisingly, we found that ∼44% of cells were able to regrow (Fig. 4C) more than 10-15 hours after infection (Fig. S4C). We wondered whether the exit from dormancy was inversely correlated with the burden of the ΦNM4γ4*^gfp^*lytic cycle for the host cell. To test this, we compared the average green fluorescence value for staphylococci that did or did not regrow in the 96-well plate after cell sorting. We found a small but significant increase in the GFP expression levels of cells that were not able to proliferate (Fig. S4D). Assuming that GFP levels are correlated with viral replication and/or expression, this result suggests that these variables impact the ability of an infected cell to leave Csm6-mediated dormancy. In contrast, the time that took each arrested cell to resume growth was variable (Fig. S4C) and did not correlate to the initial GFP intensities in the sorted cells (Fig. S4E). Sorted cells were also monitored under the microscope, and, in the absence of uninfected cells that covered the entire field of view, we were able to see staphylococci that lose GFP signal and began division. We hypothesized that the nuclease activity of Cas10, which prevents Csm6 from triggering host dormancy during *spc36* targeting and other crRNAs that target the downstream region of the PE transcript, could also enable escape from the growth arrest conveyed by guides complementary to late-expressed phage transcripts. To test this, we sorted GFP-expressing staphylococci that harbored *spc47* and the *cas10*^HD^ mutant allele (Fig. 3C) into a 96-well plate, with one cell per well, and five replicates. We found that less than 25% of cells were able to recover (Fig. 4C). Interestingly, in the absence of Cas10 nuclease activity, the initial GFP intensity did not seem to affect chances of regrowth (Fig. S4F). Overall, these data suggest that spacers that mediate the targeting of late-expressed phage transcripts are acquired and maintained in the surviving population during type III-A CRISPR-Cas immunity, in part as a result of the nuclease activity of Cas10, which presumably degrades the invading viral genome to eliminate target transcription, shut down the Csm6 response and enable a restart of bacterial growth.

### Csm6-induced dormancy provides broad-spectrum immunity

The occurrence of spacers that produce crRNAs complementary to late-expressed targets in *Staphylococcus* isolates (Cao *et al*., 2016; Wang *et al*., 2021) suggests that these spacers can confer evolutionary advantages that offset the fitness cost imposed by the growth arrest triggered by these sequences during type III-A defense. Since we previously showed that Csm6 was important to maintain immunity against phages containing mutations in their target sequence (Jiang *et al*., 2016), we hypothesized that the population immunity mediated by this RNase could protect against the rise of mutant phages with nucleotide substitutions that prevent the complete annealing of the crRNA to its target transcript and enable evasion of type III-A CRISPR immunity. To investigate this, we measured ΦNM4γ4 proliferation through the enumeration of plaque-forming units (PFU) at different times after infection of cultures carrying *spc15, spc36,* and *spc47* at an MOI of 10. Compared to a control culture of staphylococci that are unable to target the phage (Δ*spc*), which displayed a continuous PFU increase, cultures carrying *spc15* and *spc36* showed an initial decrease in the production of phage particles followed by PFU surge, approximately 2.5 hours after infection (Fig. 5A). To test for the presence of phage escapers in these cultures, we amplified and sequenced the *spc15* and *spc36* targets using DNA extracted from 10 individual plaques. We found that all of these phages contained deletions that spanned the target sequence (Table S5). To quantify the escapers, we spotted the supernatants of the liquid cultures (from the 3-hour time point in Figure 5A) on top agar containing either staphylococci harboring pCRISPR plasmids programmed with the same spacer sequence as the cells in the liquid culture, or a non-targeting host strain (*S. aureus* RN4220). We found that the supernatants of *spc15* and *spc36* cultures formed approximately the same number of plaques in both strains, a result that demonstrates that the great majority of the phages in the supernatants can escape type III-A defense (Fig. 5B). Surprisingly, in spite of the presence of *spc15* and *spc36* escapers in infected cultures, these did not affect the growth of the bacterial population (Fig. 1B) nor the viability of the staphylococci within these cultures (Fig. 1C). We hypothesized that the escaper phages reach high numbers during stationary growth, when cells are not hospitable for infection (Attrill et al., 2023). To test this possibility, we diluted *spc15* and *spc36* cultures that reached stationary phase and after treatment with ΦNM4γ4 in an MOI of 10 to an OD of 0.1. We found that the diluted cells resumed exponential growth and became susceptible to infection by phage escapers, which bypassed type III-A CRISPR immunity and caused the collapse of the population (Fig. 5C).

**Figure 5.**
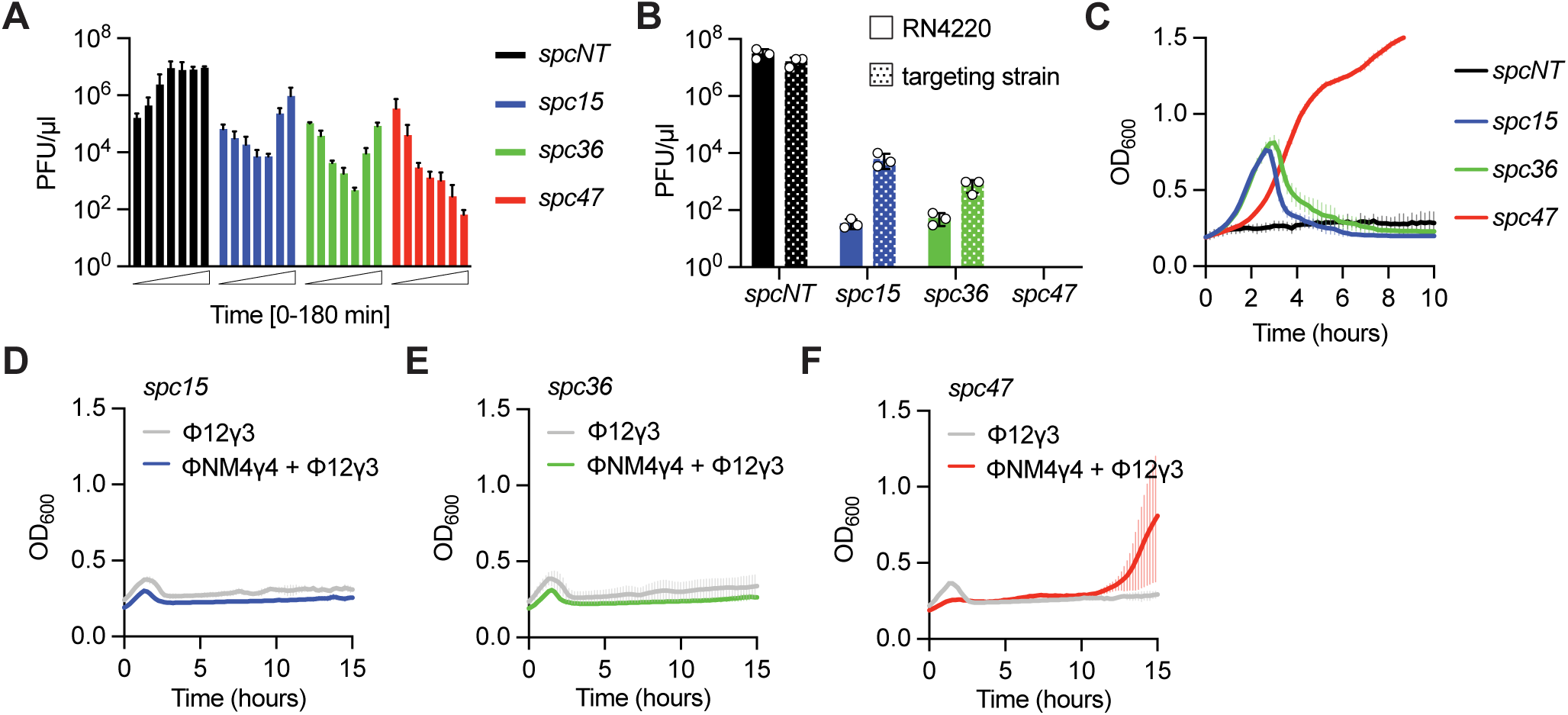
Csm6-induced dormancy provides broad-spectrum immunity. **(A)** Mean (± SD, n = 3 biological replicates) PFU/µl in supernatant of cultures harboring pCRISPR with the indicated spacers (*spc15, spc36, spc47,* Δ*spc*) samples every 30 minutes for three hours, after infection with ΦNM4γ4 at an MOI of 10. **(B)** Mean (± SD, n = 3 biological replicates) PFU/µl in supernatants of the liquid cultures shown in (A) (taken at the 3-hour time point) spotted on top agar containing either staphylococci harboring pCRISPR plasmids programmed with the same spacer sequence as the cells in the liquid culture, or a non-targeting host strain (*S. aureus* RN4220). **(C)** Mean (± SD, n = 3 biological replicates) OD_600_ values of staphylococcal cultures harboring a pCRISPR plasmid programmed with the indicated spacers (*spc15, spc36, spc47,* Δ*spc*) after infection with ΦNM4γ4 at MOI 10 for one hour in panel **(B)**, re-diluted to OD_600_ of 0.1. **(D)** Mean (± SD, n = 3 biological replicates) OD_600_ values of staphylococcal cultures harboring pCRISPR programmed with *spc15*, infected with ΦNM4γ4 at MOI 10 for one hour and then infected with Φ12γ3 at MOI 10. Infection with Φ12γ3 without prior treatment is shown as control. **(E)** Same as **(D)** but following infection of cultures programmed with *spc36*. **(F)** Same as **(E)** but following infection of cultures programmed with *spc47*.

In contrast to the detrimental rise of ΦNM4γ4 escapers in cultures harboring *spc15* or *spc36*, in agreement with the steady decrease of PFU over time in supernatants of *spc47* cultures, no escaper plaques were detected in these supernatants (Fig. 5B). In addition, the growth of infected cultures that were diluted to OD 0.1 was not affected (Fig. 5C). These results suggest that escaper phages cannot propagate in bacterial populations that, due to the activation of Csm6, contain a large fraction of dormant cells.

This is most likely due to the entry of escaper phage into dormant cells (where the type III response has been previously triggered by infection with a wild-type phage) that cannot support their propagation. If this hypothesis is correct, the cell dormancy triggered by *spc47*-mediated type III-A CRISPR immunity should not only protect against escaper phages, but also against other unrelated phages that do not have a target sequence for *spc47* crRNA. To test this, we performed sequential infection at MOI 10 of staphylococci harboring *spc15*, *spc36* or *spc47* with ΦNM4γ4 and Φ12γ3 (Modell et al., 2017), which lacks target sequences for these spacers. While *spc15* and *spc36* cultures succumbed to Φ12γ3 treatment (Figs. 5D-E), staphylococci carrying *spc47* were able survive the double infection (2/3 replicates; Figs. 5F and S5A). As expected, this cross-immunity was not observed in staphylococci expressing dCsm6 (Fig. S5B). Similar results were obtained using Φ80α*vir* (Banh *et al*., 2023) as the non-targeted phage (Figs. S5C-E). Together, these results suggest that the Csm6-induced dormancy generated during the targeting of late-expressed viral transcripts by the type III-A CRISPR-Cas system provides broad spectrum immunity against other phages present in the host environment. We believe that this feature represents a selective advantage that promotes the maintenance of spacers that mediate this type of immunity.

## DISCUSSION

Here we thoroughly characterized the Csm6 response to phage infection in staphylococci that harbor type III-A CRISPR-Cas systems. We found that Csm6 is not only required for defense when the target transcript is expressed late in the phage lytic cycle, but also when it is mediated by crRNAs that anneal to early-expressed viral RNAs. Targeting of late transcripts generated a prolonged dormancy of the infected cells that prevented phage propagation to allow the survival of uninfected cells. By isolating single infected cells in the arrested state, we found that about half of them are able to resume growth many hours after being exposed to phage. We determined that regrowth is in part enabled by the ssDNase activity of the Cas10-Csm complex. We found that dormant cells begin to replicate again approximately 15 hours after the addition of phage; on the other hand uninfected cells are able to proliferate throughout the course of infection. Therefore, majority of the bacteria that make the surviving population most likely avoided infection. For spacers that require Csm6 to provide immunity through the targeting of early-expressed viral transcripts the growth arrest was very minor and required Cas10’s ssDNase activity for its relief, presumably rapidly clearing the phage DNA. We also investigated spacer acquisition during phage infection. We found that the incorporation of spacers that trigger Csm6-mediated dormancy occurs at extremely low frequencies, a result consistent with the long time it takes for the infected cells carrying these spacers to resume growth. In contrast, spacers that produce crRNAs that recognize early-expressed transcripts and either do not depend on Csm6 for immunity, or that require Csm6 but the growth arrest is rapidly mitigated, are persistently acquired. Finally, we explored the benefits of the dormancy-based immunity that could justify an evolutionary need for the maintenance of the rarely acquired spacers that trigger it. We demonstrate that this mechanism leads to defense against not only the target phage but also other viruses present in the population that fail to replicate in the arrested cells. Altogether, our data a comprehensive model for the molecular relationship between the two activities of type III CRISPR-Cas systems. On one hand, the Cas10 complex ability to degrade viral ssDNA is sufficient to provide immunity at the individual cell level (which is rapidly “cured” from infection) when the crRNA rapidly recognizes the invading phage. On the other hand, the growth arrest mediated by CARF effectors like Csm6 confers defense at the population level, halting the propagation of both the phage and its host to allow the survival of uninfected cells. This type of immunity, however, also depends on the ssDNase activity of the type III system, which facilitates the recovery of the arrested cells that is required for the subsistence of the dormancy-triggering spacers.

An unexpected discovery of our work is the existence of a set of spacers that produce crRNAs that target the downstream region of the early-expressed ΦNM4γ4 transcript and that require the nuclease activities of both Cas10 and Csm6 for immunity. We previously hypothesized that the immunity that relies on the ssDNA degradation by the Cas10-Csm complex (which is believed to attack substrates present at transcription bubbles (Liu *et al*., 2019; Samai *et al*., 2015)) is only effective if this activity is set off almost immediately after infection, before too many copies of the viral genome are present in the host cell (Jiang *et al*., 2016). If this is the case, only crRNAs able to recognize the very first transcripts produced in the lytic cycle will mediate successful immunity, whereas crRNAs that recognize PE-derived transcripts that are produced later trigger the ssDNase activity when the viral DNA has accumulated after rounds of replication (Wegrzyn et al., 2012). In addition, the downstream region of the early-expressed ΦNM4γ4 transcript showed lower abundance of reads in our RNAseq experiments at 5 minutes post-infection (Fig. S2E). This could further decrease the efficiency of the ssDNase activity since previous data showed absence of plasmid DNA degradation by the staphylococcal type III-A CRISPR-Cas system when the target is poorly transcribed (Rostol and Marraffini, 2019), a situation that, similarly to the anti-phage immunity mediated by *spc36*, also required Csm6 for plasmid clearance.

In addition to the target region that requires both ssDNase and RNase activities for defense described above, the use of spacer libraries with high coverage density revealed other important aspects of the type III-A CRISPR response. For example, although the two modes of immunity of type III-A systems in staphylococci depend on whether the target recognized by the crRNA is an early- or late-expressed phage transcript, this classification is not absolute as there is a gradual decrease in the enrichment of spacers across the late promoter region (15 to 18 kb of the ΦNM4γ4 genome, Fig. 1A). We speculate that this is a consequence of a decrease in the ability of the ssDNase activity to clear the viral DNA, as the target transcripts are expressed at a lower level from this region (see above), which leads to a surge in the requirement of Csm6-mediated growth arrest for immunity and a lower enrichment of the cells carrying spacers that target this region. Another important finding is that, while all of the non-functional spacers that match the plus strand were present in the library population after infection of staphylococci carrying a wild-type type III-A CRISPR locus, only 20% of these spacers were detected in the presence of the d*csm6* allele (Figs. S1D and S2B-D). This result supports the conclusion that Csm6-mediated dormancy provides population-level immunity that allows the regrowth of uninfected cells. Given that these spacers cannot trigger the type III-A CRISPR response and the infection of the cells that carried them will result in lysis and death, the observation that they remain present in the population only when Csm6 is functional suggests that they are harbored by cells that avoided infection as a consequence of the dormancy triggered by this CARF effector.

We believe that the different type III immune mechanisms elicited by early- and late-transcribed targets are applicable to other phages. Indeed, this is the case for the *Sulfolobus* type III-B response against the SIRV2 and SIRV3 archaeal viruses (Lin et al., 2023; Liu et al., 2023). We believe that, although transcription of a phage genome is often not so precisely defined by two promoters as it is the case for ΦNM4γ4, most viruses have differential expression patterns to ensure the sequential completion of the different stages of the lytic cycle (Kornienko et al., 2023; Lavysh et al., 2022; Miller et al., 2003; Semenova et al., 2005; Wang et al., 2005). This is a problem for type III CRISPR immunity since the appearance of a target RNA late in infection forces a delayed response, mounted when phage replication is well under way and the host cell possibly compromised. Evolutionary evidence of this problem is the ubiquitous presence of many different CARF effectors associated with type III systems (Makarova et al., 2020). This is because, to date, all of the CARF effectors that have been characterized experimentally are capable of causing cell toxicity when activated by cOAs (Baca et al., 2024; Han et al., 2017; McMahon et al., 2020; Rostol and Marraffini, 2019; Rostol et al., 2021; Zhu et al., 2021), an observation that suggests that these mediate a mechanism of immunity that saves the population but not the compromised infected host. While CARF effectors can provide a solution to a delayed type III response, the host arrest they produce would lead to the loss of the spacers that trigger it. Our work shows that the exit from dormancy mediated by the ssDNase activity is a built-in mechanism within type III CRISPR-Cas systems to solve this second problem and allow the subsistence of spacers that provide population-level immunity.

While the above model for type III immunity is largely supported by the results in this and previous work, there are many observations that remain to be incorporated. For example, there are type III systems in which Cas10 lacks an HD domain (Makarova et al., 2015) or that do not display detectable ssDNase activity (Gruschow et al., 2019). In such instances, the exit from dormancy should rely exclusively on host nucleases or other processes that lead to the elimination of the viral genome from the arrested cell, as it is the case for the type VI-A CRISPR-Cas system of *Listeria*, which requires the activity of restriction systems to alleviate the host arrest they generate (Williams et al., 2023). In addition, many type III CRISPR loci harbor ring nucleases that degrade cOA ligands to limit the activation of CARF effectors (Athukoralage et al., 2018; Rouillon et al., 2018). How these ring nucleases modulate the type III response to affect the exit from dormancy is not known. Finally, the presence of multiple type III CRISPR loci is common among bacteria and archaea (Makarova *et al*., 2015). This adds another layer of complexity to the type III response that remains to be understood. For example, as mentioned above, the hyperthermophilic bacterium *T. thermophilus* HB27c harbors, in addition to types I-B and I-C CRISPR-Cas systems, type III-A and III-B loci with three different CARF effector genes (Lopatina et al., 2019). In this bacterium, spacers provide immunity only against targets located in an early-expressed region of the phage phiFa, and against targets transcribed across the complete genome of the phiKo phage, in both cases without producing a detectable growth arrest of host toxicity (Artamonova *et al*., 2020). Future work on hosts that naturally harbor type III loci will be needed to further our understanding of the most complex CRISPR-Cas systems.

## Methods

### Bacterial strains and growth conditions

*S. aureus* RN4220 (Kreiswirth *et al*., 1983) was incubated in brain-heart-infusion (BHI) broth at 37°C shaking (220 RPM). Cultures were supplemented with chloramphenicol (10μg/mL) to maintain CRISPR plasmids, and with spectinomycin (250 μg/mL) to maintain the spacer-library plasmids.

### Molecular cloning

Plasmids used in this study are listed in Table S6, and all oligonucleotides used are listed in Table S7. Plasmid cloning strategies are listed in Table S8.

### Spacer library construction and infection assay

For spacer library construction, a library of 40,338 × 90-nt oligonucleotides, each containing a unique ΦNM4γ4-matching spacer, type III-A repeat homology, BsaI sites, and universal priming sites, was purchased from Twist Biosciences. The library was made double-stranded by PCR with primers NA 484 and NA485, and the product was purified using MinElute PCR Purification Kit (Qiagen Cat. #28004). Spacers were introduced into pNA15 via Golden Gate cloning with BsaI-HFv2 (New England Biolabs Cat. #R3733S) and T7 DNA ligase (New England Biolabs Cat. #M0318S), then electroporated into Endura Duo electrocompetent cells (Biosearch Technologies Cat. # 60242). Five hundred thousand colonies were pooled, and the plasmids were isolated and electroporated into *S. aureus* cells harboring either pNA1, pNA23 or pNA24. Five hundred thousand CFU of each background were pooled and grown overnight. Overnight cultures were diluted 1:100 in 50 ml BHI supplemented with 5uM CaCl_2_ and appropriate antibiotics for selection. Cultures were grown shaking at 37℃ until OD_600_ of 0.2 was reached. An uninfected sample (t_0_) was obtained by pelleting 10 ml of the culture and removing supernatant. Pellets were kept at −80°C until all time points were collected. MOI of 2 of ΦNM4γ4 phage was added to before it was placed back shaking at 37°C. 5 hr post-infection 10 ml of the culture was pelleted and kept at −80°C. 24 hr post-infection 3 ml of the culture was pelleted and kept at −80°C.

### Naïve spacer acquisition assay

Overnight cultures of *S. aureus* RN4220 containing pNA2, pNA3, pNA4 or pGG32 (Heler *et al*., 2015) were diluted 1:100 in 10 mL of BHI and grown for 2.5 hr shaking at 37°C. OD_600_ was measured and used to calculate CFU per μL. For the infection, 1.3 billion CFUs were mixed with 2.6 PFUs of ΦNM4γ4 (Heler *et al*., 2015) for a starting MOI of 2. This mixture was added to 6 mL of melted 50% Heart Infusion Agar (HIA top agar) supplemented with 5 mM CaCl2. The top agar was poured onto a plate containing solidified BHI agar, and plates were then incubated at 37°C for 48 hr. To collect all colonies, top agar was scraped into a 50 ml falcon tube and incubated in a thermoblock until agar was solubilized. Cells were pelleted and supernatant was removed before quickly resuming with spacer selection (described below).

### Spacer selection and deep sequencing

Plasmids were isolated from *S. aureus* pellets with a modified protocol of QIAprep Spin Miniprep Kit (Qiagen Cat. #27104), as previously described (Aviram *et al*., 2022). 250 ng of each sample were used as input for PCR, using the Phusion DNA Polymerase (Thermo Cat. #F530L) with primers NA162/NA163 for the spacer-library experiments, and with NA101/NA102 or NA169/170 for type III-A or type II-A naïve spacer acquisition, respectively (see Table S7 for a full list of primers used in this study).

Spacer library PCRs were analyzed by agarose gel electrophoresis, bands were excised and purified using QIAquick gel extraction kit (Qiagen Cat. #28704). Naïve spacer acquisition PCRs underwent cleanup with the MinElute PCR Purification Kit (Qiagen Cat. #28004) and size selection using PippnHT 3% cassette with a timed protocol set at extraction between 26 and 35 min. Size selected products were then prepared for sequencing with the TrueSeq Nano DNA Library Prep protocol (Illumina). For maintaining the small sized product, 2.2× Sample Purification Beads (Ilumina) were used after end repair. Illumina libraries underwent high-throughput sequencing with the MiSeq platform.

### ΦNM4γ4 RNA sequencing

Overnight culture of RN4220 was diluted 1:200 in 50 ml BHI supplemented with 5mM CaCl_2_ and grown 70 minutes shaking at 37°C. OD600 was measured and MOI of 2 of ΦNM4γ4 phage was added to the culture before it was placed back shaking at 37°C. At 5, 15 and 30 minutes post-infection, 10 ml of the cultures were added to 50 ml falcon tubes containing 35 ml of ice-cold BHI and centrifuged at 4°C 120k rpm for 2 minutes. Supernatant was removed and pellets were resuspended in 100ul RNase free PBS supplemented with 100 μg/ml lysostaphin (AMBI Products), incubated for 5 min at 37°C, and sarkosyl was added at 1%. RNA was purified from lysed pellets using the Zymo Direct-Zol RNA miniprep plus kit, and genomic DNA was removed by Ambion Turbo DNA-free kit. Illumina Ribo-Zero rRNA removal (Bacteria) kit was used to remove rRNA. Samples were prepared for sequencing using the TruSeq Stranded mRNA Library prep kit, beginning at the RNA fragmentation step. Illumina libraries underwent high-throughput sequencing with the MiSeq platform.

### High-throughout sequencing data analysis

Spacers were extracted from MiSeq FASTQ files using a Python code that finds all sequences flanked by two DR sequences. For spacer-library experiment, each spacer was counted by a python code and exported to excel. Spacer frequencies at each time-point were calculated and enrichment ratios were obtained by dividing each spacer frequency at 5 hr and 24 hr time by the frequency at t_0_.

For naïve spacer-acquisition assay, genome alignment maps were generated by alignment of all spacers to the ΦNM4γ4 genome using bowtie2. Genome positions covered by aligned spacers were counted and aggregated using the Python pysam package (version 0.15.3). Reads were analyzed at a single nucleotide resolution, and RPM values were calculated as phage reads per million total aligned reads. RNA-seq sequences underwent the same pipeline of analysis, and number of reads were normalized to account for the variability in total number of reads per time point.

### Liquid culture growth assays

Overnight cultures of *S. aureus* RN4220 harboring pCRISPR were diluted 1:100 in BHI supplemented with 5uM CaCl_2_ and appropriate antibiotics for selection. Cultures were grown shaking at 37℃ for 1.5 hr and re-diluted to OD_600_ of 0.1. 200ul of each culture was seeded into each well in a 96-well plate. A calculated volume of phage stock to give an MOI of 1 or 10 was added to conditions with phage. No phage was added to control conditions. OD_600_ was measured every 10 minutes (TECAN Infinite 200 PRO) at 37 °C with shaking.

### CFU growth arrest assay

Overnight cultures of *S. aureus* RN4220 harboring pCRISPR were diluted 1:100 in BHI supplemented with 5uM CaCl_2_ and appropriate antibiotics for selection. Cultures were grown shaking at 37℃ for 1.5 hr and re-diluted to OD_600_ of 0.1. Volumes were divided between no-phage and with-phage conditions. A 1ml pre-infection aliquot was sampled from each culture. A calculated volume of phage stock to give MOI 10 was added to with-phage conditions. 1ml aliquots of all cultures were sampled every 30 minutes for 4 hours. Each aliquot was spun down at 3700 rpm for 5 minutes. The supernatant was collected and stored for further phage titer calculation. The pelleted cells were washed twice in 1ml BHI and finally resuspended in 100ul of BHI.1:10 serial dilutions were carried out. 5ul of each dilution was plated onto BHI agar plates supplemented with antibiotics for selection. Plates were incubated at 37 °C overnight. Colonies were counted the following morning to calculate total CFU/ul of plated culture.

### Microscopy time course and analysis

Overnight cultures of *S. aureus* RN4220 harboring pCRISPR were diluted 1:200 in BHI supplemented with 2.5uM CaCl_2_ and appropriate antibiotics for selection. Cells were loaded into microfluidic chambers using the CellASIC ONIX2 microfluidic system (Millipore-Sigma). After cells became trapped in the chamber, they were supplied with BHI medium under a constant flow of 5 µl /hr for 2 hours to allow them to equilibrate. Phage stock at 10^7^ concentration was added to the chamber for 20 min under a constant flow of 5 µl /hr. Flow was then switched back to BHI medium for the remainder of imaging. Phase contrast images were captured at x1000 magnification every 2 min, using a Nikon Ti2e inverted microscope equipped with a Hamamatsu Orca-Fusion SCMOS camera with the temperature-controlled enclosure set to 37 °C. GFP signal was imaged with a GFP filter set using an Excelitas Xylis LED Illuminator set to 5% power, with an exposure time of 200 ms. GFP images were captured at x1000 magnification every 10 minutes. Timelapse images were aligned and processed using NIS Elements software v5.3. The timelapse files were converted to avi and mp4 videos using a 100 ms frame rate.

### Fluorescently activated single-cell sorting

Overnight cultures of *S. aureus* RN4220 harboring pCRISPR were diluted 1:100 in BHI supplemented with 5uM CaCl_2_ and appropriate antibiotics for selection. Cultures were grown shaking at 37 °C for 1 hr, re-diluted to OD_600_ of 0.1 and infected with ΦNM4γ4-GFP phage at MOI 10. Cultures were grown shaking at 37 °C for 1 hr. Cells were centrifuged at 3700 rpm for 5 min. The supernatant was discarded, and cells were washed twice in 5 ml of 1 x dPBS. Cells were finally resuspended in 3ml of 1x dPBS. The suspension was loaded into the FACS Aria III flow cytometer (BD, USA) with a flow rate of 10,000 events per second. Bacterial cells were identified based on FSC, SSC, and fluorescence signal (GFP) using Diva software. Cells of interest were gated, and single cells were sorted into a 96-well plate containing 200ul of BHI supplemented with antibiotics. Plates were incubated overnight (TECAN Infinite 200 PRO) at 37 °C with shaking. Further analysis was done using FlowJo v10.9.0. The initial GFP intensity of each single cell was recorded using the index-sort module on the FACS Diva software and extracted with IndexSort v3.0.7 (BD FlowJo exchange). Proper biosafety measures and instrument maintenance were followed throughout the procedure to ensure accurate and contamination-free sorting.

### Total PFU quantification

100ul of an overnight culture of *S. aureus* RN4220 (containing no CRISPR plasmid) was added to 10ml of BHI top agar supplemented with 5uM CaCl_2_ and plated on BHI agar plates. The supernatants collected from aliquots in the CFU growth arrest assay described above were filtered with 0.45um *Supor*® Membrane filters (VWR, Cat# 28143-352). 1:10 serial dilutions were carried out and 2ul of each dilution was plated onto the top agar containing RN4220. Plates were incubated at 37 °C overnight. Plaques were counted the following morning to calculate total PFU/ul of plated supernatant.

### Quantification and characterization of escaper-phages

100ul of an overnight culture containing pCRISPR was added to 10ml of BHI top agar supplemented with 5uM CaCl_2_ and the appropriate antibiotic for selection and plated on BHI agar plates. 2ul of each dilution from the phage propagation assay described above (Total PFU quantification) was plated onto the top agar containing a given pCRISPR. Plates were incubated at 37 °C overnight. Plaques were counted the following morning to calculate escaper PFU/ul. In order to purify escaper-phage plaques, 10 escaper plaques from each pCRISPR were isolated and resuspended in 30ul BHI. 1:10 serial dilutions were carried out and 2ul of the resuspension was plated again on BHI top agar containing pCRISPR. Plates were incubated at 37 °C overnight. Escaper-phage plaques were selected the following day, resuspended in 30ul of BHI and heated to 98 °C for 10 mins. PCR was carried out across spacer targets on the phage genome and sent for Sanger sequencing to determine the sequence of escaper-phages.

### Liquid culture growth assays with escaper-phages

Overnight cultures of *S. aureus* RN4220 harboring pCRISPR were diluted 1:100 in BHI supplemented with 5uM CaCl_2_ and appropriate antibiotics for selection. Cultures were grown shaking at 37 °C for 1.5 hrs and re-diluted to OD_600_ of 0.2. A calculated volume of phage stock to give an MOI 10 was added and cultures were incubated for another 1 hr at 37℃ shaking. Cultures were re-diluted to OD_600_ of 0.1 and 200ul of each culture was seeded into each well in a 96-well plate. OD_600_ was measured every 10 minutes (TECAN Infinite 200 PRO) at 37 °C with shaking.

### Sequential phage infections

Overnight cultures of *S. aureus* RN4220 harboring pCRISPR were diluted 1:100 in BHI supplemented with 5uM CaCl_2_ and appropriate antibiotics for selection. Cultures were grown shaking at 37 °C for 1.5 hrs and re-diluted to OD_600_ of 0.2. The calculated volume of ΦNM4γ4 phage to give MOI 10 was then added to conditions that required prior activation with a targeted phage. No phage was added to cultures that did not have prior infection. Cultures were incubated shaking at 37°C for 1 hour and back diluted to OD_600_ of 0.1. 200ul of each culture was seeded into each well in a 96-well plate. A calculated volume of non-targeted Φ12γ3 or Φ80α phage stock was then added to wells to give an MOI of 1 or 10. OD_600_ was measured every 10 minutes (TECAN Infinite 200 PRO) at 37 °C with shaking.

## Author contributions

NA, AKS, and LAM designed and conceived this study. NA and AKS performed all experiments, except single-cell sorting, which was performed by BSR. NGL designed, generated and tested various plasmids used in this study. AB helped with spacer acquisition experiments. NA, AKS, and LAM wrote the manuscript.

## Acknowledgements

We would like to thank all past and present Marraffini lab members for thoughtful discussions and insight. We would like to especially thank Nora Pyenson for plasmid pNP54, Maj Brodmann for plasmid pFNMB0-mCherry2, Alex Meeske for help with designing the spacer library and with microscopy, and Pascal Maguin for assistance in the creation of ΦNM4γ4-GFP. We would also like to thank Daniel Mucida for use of his fluoresently-activated cell sorter. This work was supported by a grant from the Simons Foundation (578759, Aviram), and the National Institute of Health Director’s Pioneer Award 1DP1GM128184-01 to LAM. LAM is an investigator of the Howard Hughes Medical Institute.

---

**Figure S1.**
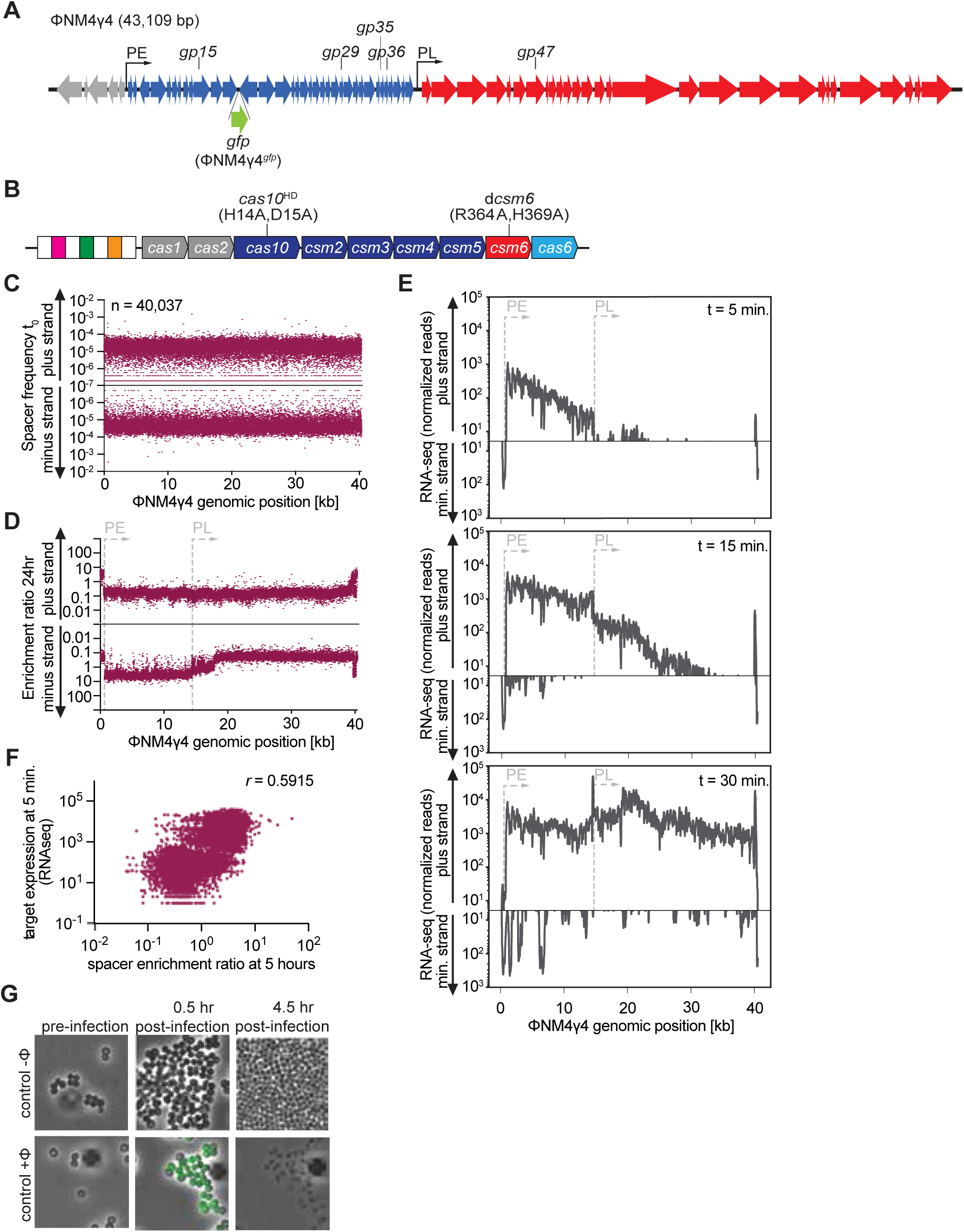
Targeting of the ΦNM4γ4 phage by the *S. epidermidis* type III-A CRISPR-Cas system. **(A)** Genome of the ΦNM4γ4 phage. Arrows indicate the two promoter, PE and PL, that drive transcription of early- and late-expressed genes. The location of the targets of the different spacers used in this study is shown. The site of insertion of *gfp* to generate ΦNM4γ4*^gfp^* is also indicated. **(B)** *S. epidermidis* type III-A CRISPR-*cas* locus. White boxes, repeats; colored boxes, spacers. Mutations introduced into *cas10* and *csm6* in this study are shown. **(C)** Spacer frequency, calculated as the fraction of spacer reads, for the library of spacers cloned into pCRISPR. Spacer sequences matching the plus and minus strands of the ΦNM4γ4 genome are plotted separately. **(D)** Enrichment ratio of spacers targeting the plus or minus strands of the ΦNM4γ4 DNA 24 hours after phage infection of staphylococci carrying pCRISPR, plotted according to their genomic position. **(E)** Abundance (in normalized reads) of ΦNM4γ4 RNA-seq reads, obtained at 5, 15 or 30 minutes after infection, mapped to the viral genome. **(F)** Correlation of target transcript expression (5 minutes after infection; shown as normalized RNA-seq reads) and enrichment of the corresponding targeting spacer (5 hours after infection). Pearson *r* coefficient is shown. **(G)** Time-course fluorescence microscopy of staphylococci at 0, 60 and 360 minutes after infection with ΦNM4γ4^GFP^ or uninfected.

**Figure S2.**
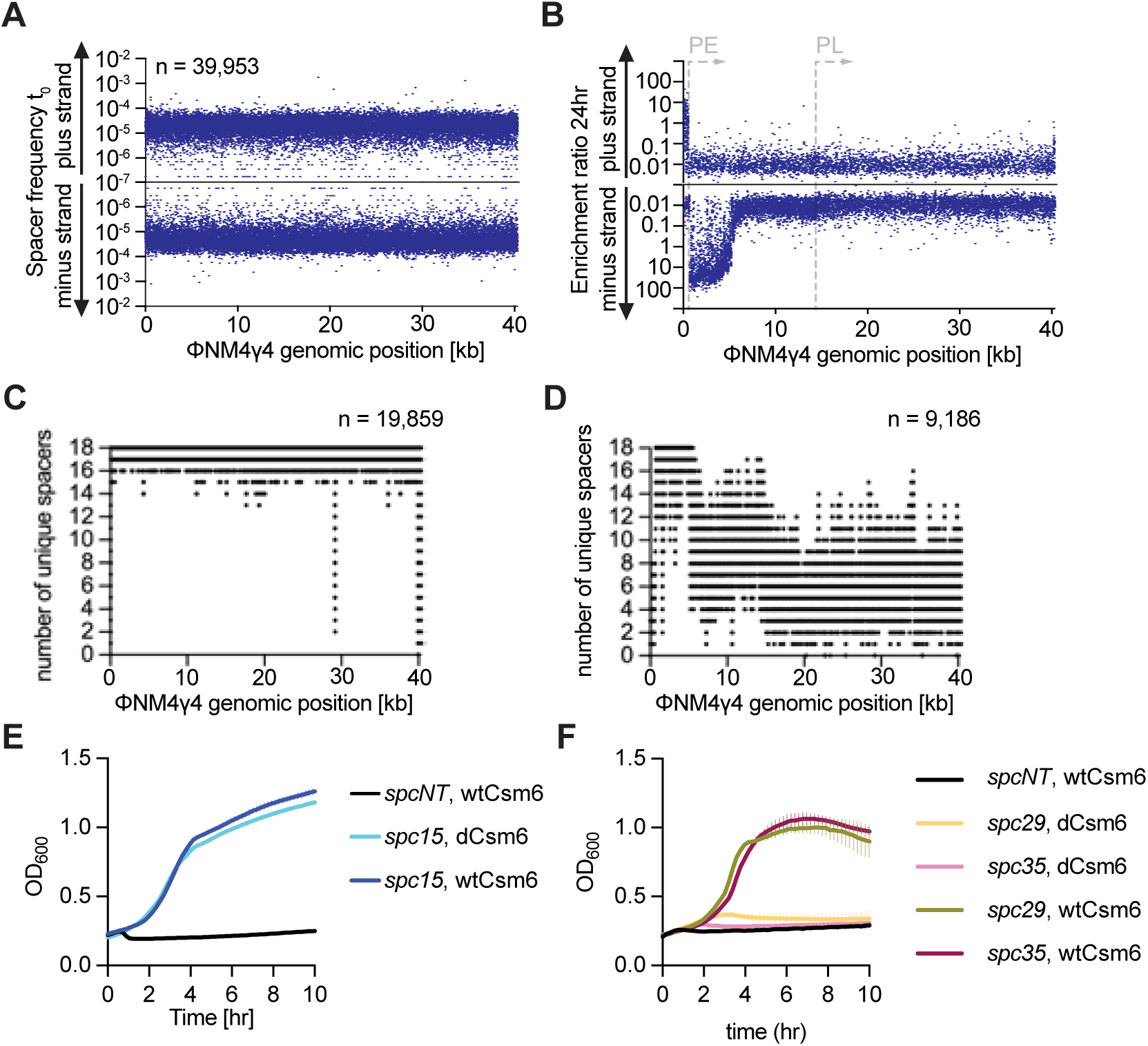
Targeting of the ΦNM4γ4 phage by a *S. epidermidis* type III-A CRISPR-Cas system carrying the d*csm6* allele. **(A)** Spacer frequency, calculated as the fraction of spacer reads, for the library of spacers cloned into pCRISPR(d*csm6*). Spacer sequences matching the plus and minus strands of the ΦNM4γ4 genome are plotted separately. **(B)** Enrichment ratio of spacers targeting the plus or minus strands of the ΦNM4γ4 DNA 24 hours after phage infection of staphylococci carrying pCRISPR(d*csm6*), plotted according to their genomic position. **(C)** Number of unique spacers detected after NGS of the pCRISPR library infected with ΦNM4γ4 for 24 hours, plotted across the viral genome. **(D)** Number of unique spacers detected after NGS of the pCRISPR(d*csm6*) library infected with ΦNM4γ4 for 24 hours, plotted across the viral genome. **(E)** Mean (± SD, n = 3 biological replicates) OD_600_ values of staphylococcal cultures harboring a pCRISPR or a pCRISPR(d*csm6*) plasmid programmed with *spc15,* after infection with ΦNM4γ4 at MOI 10. Infection of control strain programed with an non-targeting spacer is shown. **(F)** Mean (± SD, n = 3 biological replicates) OD_600_ values of staphylococcal cultures harboring a pCRISPR or a pCRISPR(d*csm6*) plasmid programmed with *spc29* or *spc35* after infection with ΦNM4γ4 at MOI 10. Infection of control strain programed with an non-targeting spacer is shown.

**Figure S3.**
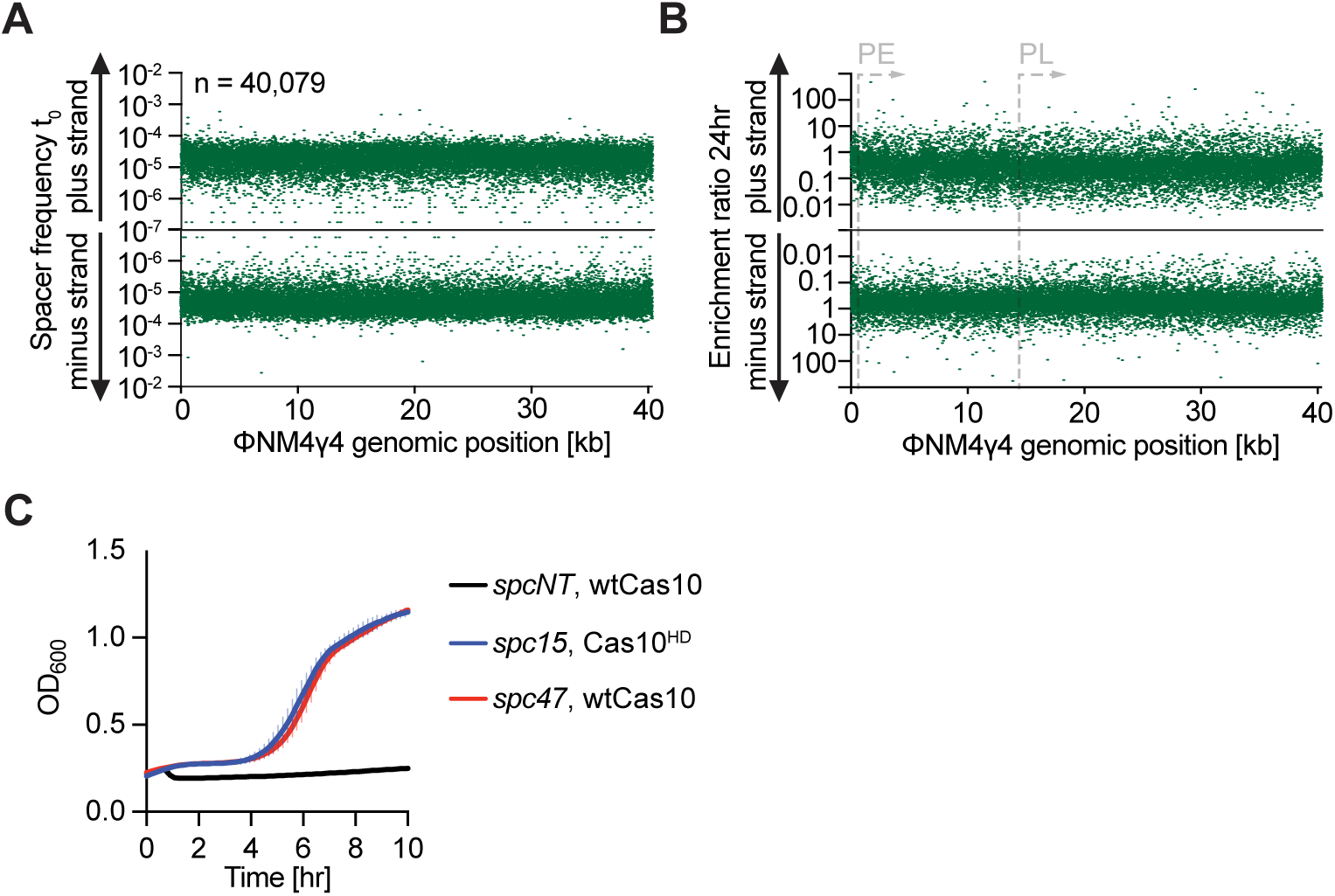
Targeting of the ΦNM4γ4 phage by a *S. epidermidis* type III-A CRISPR-Cas system carrying the *cas10*^HD^ allele. **(A)** Spacer frequency, calculated as the fraction of spacer reads, for the library of spacers cloned into pCRISPR(*cas10*^HD^). Spacer sequences matching the plus and minus strands of the ΦNM4γ4 genome are plotted separately. **(B)** Enrichment ratio of spacers targeting the plus or minus strands of the ΦNM4γ4 DNA 24 hours after phage infection of staphylococci carrying pCRISPR(*cas10*^HD^), plotted according to their genomic position. **(C)** Mean (± SD, n = 3 biological replicates) OD_600_ values of staphylococcal cultures harboring a pCRISPR or a pCRISPR(*cas10*^HD^) plasmid programmed with *spc47* or *spc15*, respectively, after infection with ΦNM4γ4 at MOI 10. Infection of control strain programed with an non-targeting spacer is shown.

**Figure S4.**
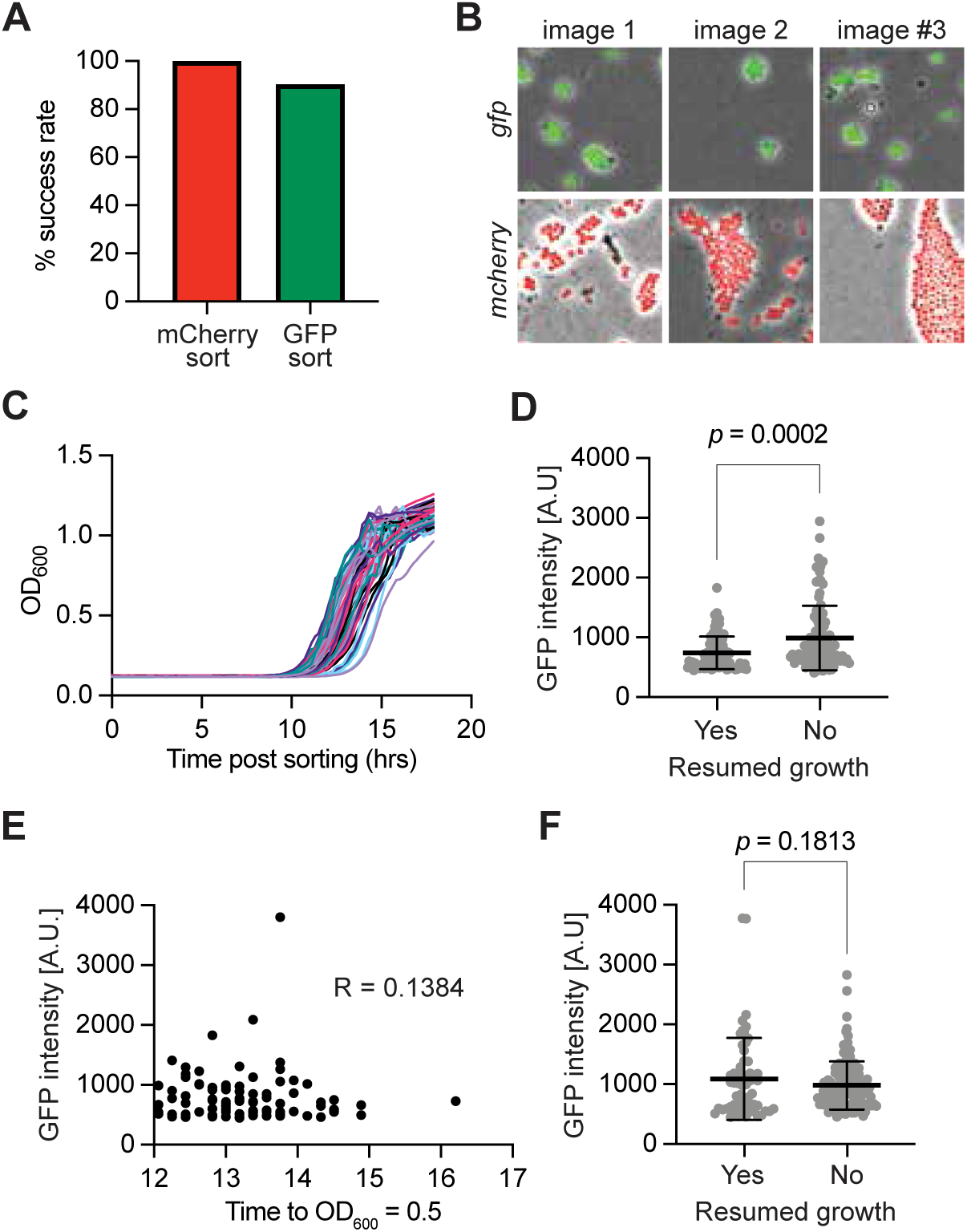
Evaluation of the exit from dormancy of infected cells expressing GFP using single-cell analyses. **(A)** Single-cell sorting control. A mixture of cells expressing mCherry or GFP were sorted into each well of a 96 well plate using gating strategies for either mCherry or GFP. A successful sort was classified as a well that only contained the fluorescent protein that was sorted for, which was determined by examining the cells under the microscope (see panel **(B)**). The percentage of successful sorts compared to the total number of wells sorted for each is plotted. **(B)** Images of GFP- and mCherry-sorted cells. Three representative images are shown for each condition. **(C)** Growth of sorted single cells (harboring pCRISPR programmed with *spc47* and infected with ΦNM4γ4*^gfp^*) within a 96-well plate measured as OD_600_ values over time after sorting. **(D)** Mean (± SD, n=2 independent experiments) initial GFP intensity of ΦNM4γ4*^gfp^*-infected staphylococci harboring pCRISPR programmed with *spc47* during indexed cell sorting, categorized based on growth outcome (*p* value obtained by unpaired t-test). **(E)** Correlation of initial GFP intensity and the time (hours) a given well reached OD_600_ of 0.5. Pearson *R* coefficient is depicted on the plot. (F) Same as (D) but using staphylococci harboring pCRISPR(*cas10*^HD^) programmed with *spc47*.

**Figure S5.**
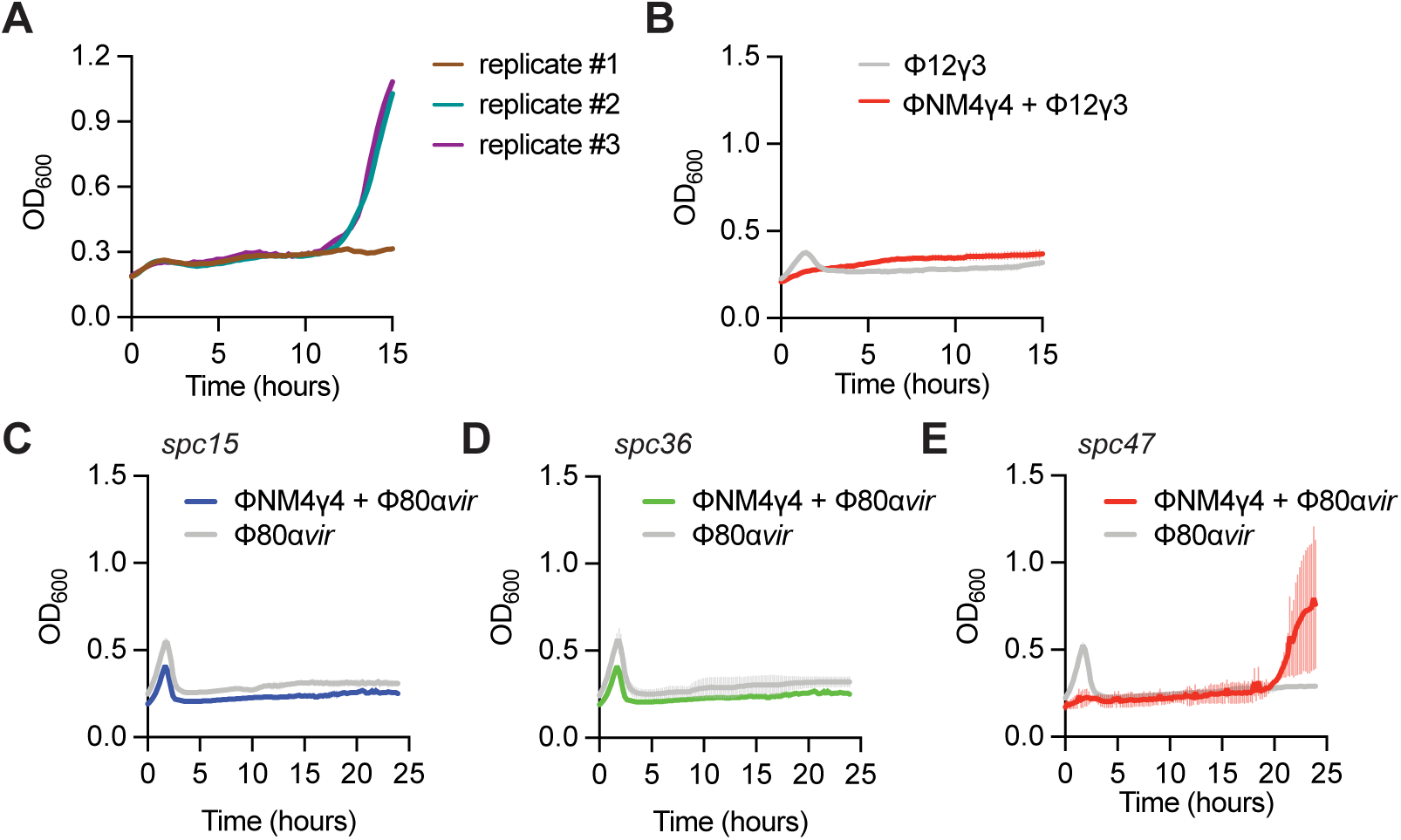
Type III-A immunity protects staphylococci against mixed pahge infections. **(A)** Growth of individual culture replicates shown in Figure 5F. **(B)** Mean (± SD, n = 3 biological replicates) OD_600_ values of staphylococcal cultures harboring pCRISPR(d*csm6*) programmed with *spc47*, infected with ΦNM4γ4 at MOI 10 for one hour and then infected with Φ12γ3 at MOI 10. Infection with Φ12γ3 without prior treatment is shown as control. **(C)** Mean (± SD, n = 3 biological replicates) OD_600_ values of staphylococcal cultures harboring pCRISPR programmed with *spc15*, infected with ΦNM4γ4 at MOI 10 for one hour and then infected with Φ80α*vir* at MOI 1. Infection with Φ80α*vir* without prior treatment is shown as control. **(D)** Same as **(C)** but following infection of cultures programmed with *spc36*. **(E)** Same as **(D)** but following infection of cultures programmed with *spc47*.

## REFERENCES

Abudayyeh, O.O., Gootenberg, J.S., Essletzbichler, P., Han, S., Joung, J., Belanto, J.J., Verdine, V., Cox, D.B.T., Kellner, M.J., Regev, A., et al. (2017). RNA targeting with CRISPR-Cas13. Nature 550, 280–284.

Artamonova, D., Karneyeva, K., Medvedeva, S., Klimuk, E., Kolesnik, M., Yasinskaya, A., Samolygo, A., and Severinov, K. (2020). Spacer acquisition by Type III CRISPR-Cas system during bacteriophage infection of Thermus thermophilus. Nucleic Acids Res. 48, 9787–9803.

Athukoralage, J.S., Rouillon, C., Graham, S., Gruschow, S., and White, M.F. (2018). Ring nucleases deactivate type III CRISPR ribonucleases by degrading cyclic oligoadenylate. Nature 562, 277–280.

Attrill, E.L., Lapinska, U., Westra, E.R., Harding, S.V., and Pagliara, S. (2023). Slow growing bacteria survive bacteriophage in isolation. ISME Commun 3, 95.

Aviram, N., Thornal, A.N., Zeevi, D., and Marraffini, L.A. (2022). Different modes of spacer acquisition by the *Staphylococcus epidermidis* type III-A CRISPR-Cas system. Nucleic Acids Res. 50, 1661–1672.

Baca, C.F., Yu, Y., Rostol, J.T., Majumder, P., Patel, D.J., and Marraffini, L.A. (2024). The CRISPR effector Cam1 mediates membrane depolarization for phage defence. Nature 625, 797–804.

Bae, T., Baba, T., Hiramatsu, K., and Schneewind, O. (2006). Prophages of *Staphylococcus aureus* Newman and their contribution to virulence. Mol. Microbiol. 62, 1035–1047.

Banh, D.V., Roberts, C.G., Morales-Amador, A., Berryhill, B.A., Chaudhry, W., Levin, B.R., Brady, S.F., and Marraffini, L.A. (2023). Bacterial cGAS senses a viral RNA to initiate immunity. Nature 623, 1001–1008.

Barrangou, R., Fremaux, C., Deveau, H., Richards, M., Boyaval, P., Moineau, S., Romero, D.A., and Horvath, P. (2007). CRISPR provides acquired resistance against viruses in prokaryotes. Science 315, 1709–1712.

Bolotin, A., Quinquis, B., Sorokin, A., and Ehrlich, S.D. (2005). Clustered regularly interspaced short palindrome repeats (CRISPRs) have spacers of extrachromosomal origin. Microbiology 151, 2551–2561.

Cao, L., Gao, C.H., Zhu, J., Zhao, L., Wu, Q., Li, M., and Sun, B. (2016). Identification and functional study of type III-A CRISPR-Cas systems in clinical isolates of Staphylococcus aureus. Int. J. Med. Microbiol. 306, 686–696.

Elmore, J.R., Sheppard, N.F., Ramia, N., Deighan, T., Li, H., Terns, R.M., and Terns, M.P. (2016). Bipartite recognition of target RNAs activates DNA cleavage by the Type III-B CRISPR-Cas system. Genes Dev. 30, 447–459.

Estrella, M.A., Kuo, F.T., and Bailey, S. (2016). RNA-activated DNA cleavage by the Type III-B CRISPR-Cas effector complex. Genes Dev. 30, 460–470.

Goldberg, G.W., Jiang, W., Bikard, D., and Marraffini, L.A. (2014). Conditional tolerance of temperate phages via transcription-dependent CRISPR-Cas targeting. Nature 514, 633–637.

Gruschow, S., Athukoralage, J.S., Graham, S., Hoogeboom, T., and White, M.F. (2019). Cyclic oligoadenylate signalling mediates Mycobacterium tuberculosis CRISPR defence. Nucleic Acids Res. 47, 9259–9270.

Hale, C.R., Zhao, P., Olson, S., Duff, M.O., Graveley, B.R., Wells, L., Terns, R.M., and Terns, M.P. (2009). RNA-guided RNA cleavage by a CRISPR RNA-Cas protein complex. Cell 139, 945–956.

Han, W., Li, Y., Deng, L., Feng, M., Peng, W., Hallstrom, S., Zhang, J., Peng, N., Liang, Y.X., White, M.F., and She, Q. (2017). A type III-B CRISPR-Cas effector complex mediating massive target DNA destruction. Nucleic Acids Res. 45, 1983–1993.

Hatoum-Aslan, A., Samai, P., Maniv, I., Jiang, W., and Marraffini, L.A. (2013). A ruler protein in a complex for antiviral defense determines the length of small interfering CRISPR RNAs. J. Biol. Chem. 288, 27888–27897.

Heler, R., Samai, P., Modell, J.W., Weiner, C., Goldberg, G.W., Bikard, D., and Marraffini, L.A. (2015). Cas9 specifies functional viral targets during CRISPR-Cas adaptation. Nature 519, 199–202.

Horinouchi, S., and Weisblum, B. (1982). Nucleotide sequence and functional map of pC194, a plasmid that specifies inducible chloramphenicol resistance. J. Bacteriol. 150, 815–825.

Jiang, W., Samai, P., and Marraffini, L.A. (2016). Degradation of phage transcripts by CRISPR-associated RNases enables type III CRISPR-Cas immunity. Cell 164, 710–721.

Kazlauskiene, M., Kostiuk, G., Venclovas, C., Tamulaitis, G., and Siksnys, V. (2017). A cyclic oligonucleotide signaling pathway in type III CRISPR-Cas systems. Science 357, 605–609.

Kazlauskiene, M., Tamulaitis, G., Kostiuk, G., Venclovas, C., and Siksnys, V. (2016). Spatiotemporal Control of Type III-A CRISPR-Cas Immunity: Coupling DNA Degradation with the Target RNA Recognition. Mol. Cell 62, 295–306.

Kenney, C.T., and Marraffini, L.A. (2023). Rarely acquired type II-A CRISPR-Cas spacers mediate anti-viral immunity through the targeting of a non-canonical PAM sequence. Nucleic Acids Res. 51, 7438–7450.

Koonin, E.V., and Makarova, K.S. (2022). Evolutionary plasticity and functional versatility of CRISPR systems. PLoS Biol. 20, e3001481.

Kornienko, M., Bespiatykh, D., Gorodnichev, R., Abdraimova, N., and Shitikov, E. (2023). Transcriptional Landscapes of Herelleviridae Bacteriophages and Staphylococcus aureus during Phage Infection: An Overview. Viruses 15.

Kreiswirth, B.N., Lofdahl, S., Betley, M.J., O’Reilly, M., Schlievert, P.M., Bergdoll, M.S., and Novick, R.P. (1983). The toxic shock syndrome exotoxin structural gene is not detectably transmitted by a prophage. Nature 305, 709–712.

Lavysh, D., Mekler, V., Klimuk, E., and Severinov, K. (2022). Regulation of Gene Expression of phiEco32-like Bacteriophage 7-11. Viruses 14.

Lin, J., Alfastsen, L., Bhoobalan-Chitty, Y., and Peng, X. (2023). Molecular basis for inhibition of type III-B CRISPR-Cas by an archaeal viral anti-CRISPR protein. Cell Host Microbe 31, 1837–1849 e1835.

Liu, J., Li, Q., Wang, X., Liu, Z., Ye, Q., Liu, T., Pan, S., and Peng, N. (2023). An archaeal virus-encoded anti-CRISPR protein inhibits type III-B immunity by inhibiting Cas RNP complex turnover. Nucleic Acids Res. 51, 11783–11796.

Liu, T.Y., Liu, J.J., Aditham, A.J., Nogales, E., and Doudna, J.A. (2019). Target preference of Type III-A CRISPR-Cas complexes at the transcription bubble. Nat Commun 10, 3001.

Lopatina, A., Medvedeva, S., Artamonova, D., Kolesnik, M., Sitnik, V., Ispolatov, Y., and Severinov, K. (2019). Natural diversity of CRISPR spacers of Thermus: evidence of local spacer acquisition and global spacer exchange. Philos Trans R Soc Lond B Biol Sci 374, 20180092.

Makarova, K.S., Timinskas, A., Wolf, Y.I., Gussow, A.B., Siksnys, V., Venclovas, C., and Koonin, E.V. (2020). Evolutionary and functional classification of the CARF domain superfamily, key sensors in prokaryotic antivirus defense. Nucleic Acids Res. 48, 8828–8847.

Makarova, K.S., Wolf, Y.I., Alkhnbashi, O.S., Costa, F., Shah, S.A., Saunders, S.J., Barrangou, R., Brouns, S.J., Charpentier, E., Haft, D.H., et al. (2015). An updated evolutionary classification of CRISPR-Cas systems. Nat. Rev. Microbiol. 13, 722–736.

McMahon, S.A., Zhu, W., Graham, S., Rambo, R., White, M.F., and Gloster, T.M. (2020). Structure and mechanism of a Type III CRISPR defence DNA nuclease activated by cyclic oligoadenylate. Nat Commun 11, 500.

Meeske, A.J., Nakandakari-Higa, S., and Marraffini, L.A. (2019). Cas13-induced cellular dormancy prevents the rise of CRISPR-resistant bacteriophage. Nature 570, 241–245.

Miller, E.S., Kutter, E., Mosig, G., Arisaka, F., Kunisawa, T., and Ruger, W. (2003). Bacteriophage T4 genome. Microbiol. Mol. Biol. Rev. 67, 86–156, table of contents.

Modell, J.W., Jiang, W., and Marraffini, L.A. (2017). CRISPR-Cas systems exploit viral DNA injection to establish and maintain adaptive immunity. Nature 544, 101–104.

Mojica, F.J., Diez-Villasenor, C., Garcia-Martinez, J., and Soria, E. (2005). Intervening sequences of regularly spaced prokaryotic repeats derive from foreign genetic elements. J. Mol. Evol. 60, 174–182.

Niewoehner, O., Garcia-Doval, C., Rostol, J.T., Berk, C., Schwede, F., Bigler, L., Hall, J., Marraffini, L.A., and Jinek, M. (2017). Type III CRISPR-Cas systems produce cyclic oligoadenylate second messengers. Nature 548, 543–548.

Niewoehner, O., and Jinek, M. (2016). Structural basis for the endoribonuclease activity of the type III-A CRISPR-associated protein Csm6. RNA 22, 318–329.

Nunez, J.K., Lee, A.S., Engelman, A., and Doudna, J.A. (2015). Integrase-mediated spacer acquisition during CRISPR-Cas adaptive immunity. Nature 519, 193–198.

Perez-Casal, J., Caparon, M.G., and Scott, J.R. (1991). Mry, a trans-acting positive regulator of the M protein gene of *Streptococcus pyogenes* with similarity to the receptor proteins of two-component regulatory systems. J. Bacteriol. 173, 2617–2624.

Pourcel, C., Salvignol, G., and Vergnaud, G. (2005). CRISPR elements in *Yersinia pestis* acquire new repeats by preferential uptake of bacteriophage DNA, and provide additional tools for evolutionary studies. Microbiology 151, 653–663.

Pyenson, N.C., Gayvert, K., Varble, A., Elemento, O., and Marraffini, L.A. (2017). Broad Targeting Specificity during Bacterial Type III CRISPR-Cas Immunity Constrains Viral Escape. Cell Host Microbe 22, 343–353 e343.

Pyenson, N.C., and Marraffini, L.A. (2020). Co-evolution within structured bacterial communities results in multiple expansion of CRISPR loci and enhanced immunity. Elife 9.

Rostol, J.T., and Marraffini, L.A. (2019). Non-specific degradation of transcripts promotes plasmid clearance during type III-A CRISPR-Cas immunity. Nat Microbiol 4, 656–662.

Rostol, J.T., Xie, W., Kuryavyi, V., Maguin, P., Kao, K., Froom, R., Patel, D.J., and Marraffini, L.A. (2021). The Card1 nuclease provides defence during type III CRISPR immunity. Nature 590, 624–629.

Rouillon, C., Athukoralage, J.S., Graham, S., Gruschow, S., and White, M.F. (2018). Control of cyclic oligoadenylate synthesis in a type III CRISPR system. Elife 7.

Samai, P., Pyenson, N., Jiang, W., Goldberg, G.W., Hatoum-Aslan, A., and Marraffini, L.A. (2015). Co-transcriptional DNA and RNA Cleavage during Type III CRISPR-Cas Immunity. Cell 161, 1164–1174.

Semenova, E., Djordjevic, M., Shraiman, B., and Severinov, K. (2005). The tale of two RNA polymerases: transcription profiling and gene expression strategy of bacteriophage Xp10. Mol. Microbiol. 55, 764–777.

Sheppard, N.F., Glover, C.V., 3rd, Terns, R.M., and Terns, M.P. (2016). The CRISPR-associated Csx1 protein of Pyrococcus furiosus is an adenosine-specific endoribonuclease. RNA 22, 216–224.

Stella, G., and Marraffini, L. (2024). Type III CRISPR-Cas: beyond the Cas10 effector complex. Trends Biochem. Sci. 49, 28–37.

Wang, J., Jiang, Y., Vincent, M., Sun, Y., Yu, H., Wang, J., Bao, Q., Kong, H., and Hu, S. (2005). Complete genome sequence of bacteriophage T5. Virology 332, 45–65.

Wang, Y., Mao, T., Li, Y., Xiao, W., Liang, X., Duan, G., and Yang, H. (2021). Characterization of 67 Confirmed Clustered Regularly Interspaced Short Palindromic Repeats Loci in 52 Strains of Staphylococci. Front Microbiol 12, 736565.

Wegrzyn, G., Licznerska, K., and Wegrzyn, A. (2012). Phage lambda--new insights into regulatory circuits. Adv. Virus Res. 82, 155–178.

Williams, M.C., Reker, A.E., Margolis, S.R., Liao, J., Wiedmann, M., Rojas, E.R., and Meeske, A.J. (2023). Restriction endonuclease cleavage of phage DNA enables resuscitation from Cas13-induced bacterial dormancy. Nat Microbiol 8, 400–409.

Yosef, I., Goren, M.G., and Qimron, U. (2012). Proteins and DNA elements essential for the CRISPR adaptation process in *Escherichia coli*. Nucleic Acids Res. 40, 5569–5576.

Zhang, X., Garrett, S., Graveley, B.R., and Terns, M.P. (2021). Unique properties of spacer acquisition by the type III-A CRISPR-Cas system. Nucleic Acids Res.

Zhu, W., McQuarrie, S., Gruschow, S., McMahon, S.A., Graham, S., Gloster, T.M., and White, M.F. (2021). The CRISPR ancillary effector Can2 is a dual-specificity nuclease potentiating type III CRISPR defence. Nucleic Acids Res. 49, 2777–2789.

