## Supplementary material for "The Cas10 nuclease activity relieves host dormancy to facilitate spacer acquisition and retention during type III-A CRISPR immunity": Table S4

| **Spacer sequence** | **Staphylococcal**  **phage** | **Target gene** | **Exp.*** |
| --- | --- | --- | --- |
| GAGAACTTAATTGCATTATCAAATGTATATGCTGGATTCCA | phi66 | SPV66_ORF017 | late |
| GAGAACCCGAATTTTGATTCTTTGTTTGTAAATAATGCTC | SAP-2 | SAP2_gp03 | early |
| GAGAACCACGCTGTAGTGAAGTATAGAAACGGCATGAGTACAA | 52A | ST52AORF079 | late |
| AGTCAATATAAAGACAATACTTTTTACGCTTATATT | phiIBB-SEP1 | FDH45_gp044 | NA |
| AATGAAATTTATCAAAACTATAGAAAACTTATTAG | vB_Sau_CG | vBSauCG_155 | late |
| TGGTTTAAGTTTGTCATTATAATCAATCCTTTTTCTT | pSa-3 | pSa3_007 | early |
| GTTTTTCATAGTTAATCAATCCCTTTTCTTTTTT | pSa-3 | pSa3_067 | NA |
| TTAAATCTTTGATTGCTCTTAGCTCTAGTTATGTAT | Bacterio88 | ST88ORF056 | late |
| CACGCTGTAGTGAAGTATAGAAACGGCATGAGTACAAT | 52A | ST52AORF079 | late |
| CTTCCGAATCCATTTCAGCGCAATAAACA | SpT99F3 | SpT99F3_045 | NA |
| ACTCACTTGTAAATTCCTCCACTTGCTCTA | Bacterio2638A | ST2638AORF019 | late |
| CTGGAATAACCACAAAGCCAGAGTCAGTTT | phi575 | NA | NA |
| ACGTTAGATTTGCAGGTGTTAAGCACGGCT | vB_SepS_SEP9 | SEP9_018 | NA |
| TAGAATGTTATTATCTAAGTGGTCGATGTATTCC | BacterioG1 | ORF135-ORF092 | late |
| TTTTCTTTAACTGTTTTTACTGCCCATTTAATAGT | phiIPLA-C1C | AVU40_gp150 | late |
| AAGTTAACGGCATTACCTAATAAAAATATTTTAGG | SAP-2 | SAP2_gp08 | early |
| GTTTTTCATAGTTAATCAATCCCTTTTCTTTTTT | vB_SauM_Remus | O151_gp047 | NA |
| TATGTATTGATCTCGATTCTCGTTAGTTTCTAAATT | phiNM1 | SAPPV1_gp32 | early |
| CACGCTGTAGTGAAGTATAGAAACGGCATGAGTACAAT | SP6 | SP6_0011 | early |
| TAGTAATAATTGTCTCATTTGCATACGTTACATCGAT | CNPH82 | cn20 | NA |
| TAGAATGTTATTATCTAAGTGGTCGATGTATTCC | S25-3 | X600_gp192 | early |
| AAGTTAACGGCATTACCTAATAAAAATATTTTAGG | S13' | ORF9 | early |
| CCAAACCATTTAGCACGATATTTATTAAAACCATA | K | gp011 | early |
| TATTTTTCTCCTTTAGCAATCATTCTGTCTAGTAC | S25-3 | X600_gp187 | early |
| * Time of expression was inferred from the position on the target gene in the viral genome. NA: not assigned (difficult to determine expression timing). | | | |

**Table S4. Spacers found in staphylococcal type III-A CRISPR-Cas systems that match staphylococcal annotated phages**
