## Supplementary material for "The Cas10 nuclease activity relieves host dormancy to facilitate spacer acquisition and retention during type III-A CRISPR immunity": Table S5

**Table S5. Escapers sequenced in this study.**

| **Spacer** | **Target location (*)** | **Target deletion (*)** | **Deletion size (bp)** | **# of escapers** |
| --- | --- | --- | --- | --- |
| *spc15* | 4,236…4,270 | 3,807…5,055 | 1,249 | 2/10 |
| *spc15* | 4,236…4,270 | 3,807…5,571 | 1,765 | 7/10 |
| *spc15* | 4,236…4,270 | 3,807...5,693 | 1,887 | 1/10 |
| *spc36* | 12,909…12,943 | 12,930...12,932 | 3 | 4/10 |
| *spc36* | 12,909…12,943 | 12,926…13,015 | 90 | 1/10 |
| *spc36* | 12,909…12,943 | 12,931…13,023 | 93 | 2/10 |
| *spc36* | 12,909…12,943 | 12,894…12,943 | 50 | 2/10 |
| *spc36* | 12,909…12,943 | 12,807…13,126 | 320 | 1/10 |

(*) location within the genomic coordinates of the ΦNM4γ4 phage (GenBank accession KP209285.1)
