## Supplementary material for "The Cas10 nuclease activity relieves host dormancy to facilitate spacer acquisition and retention during type III-A CRISPR immunity": Table S6

| **Plasmid name** | **Description** | **Reference** |
| --- | --- | --- |
| pNP54 | pCRISPR *S. epidermidis* type III-A *spc47.* Used to clone pAS04, pAS55 | (2) |
| pCN57 | GFP expression. Used to clone pAS23 | (3) |
| pAS04 | pCRISPR *S. epidermidis* type III-A *spc47:dcsm6* | This study |
| pAS20 | pCRISPR *S. epidermidis* type III-A *spc15.* Used to clone pAS21, pAS53 | This study |
| pAS21 | pCRISPR *S. epidermidis* type III-A *spc15:dcsm6* | This study |
| pAS22 | Used to clone ΦNM4γ4-GFP | This study |
| pAS23 | Used to clone ΦNM4γ4-GFP | This study |
| pAS53 | pCRISPR *S. epidermidis* type III-A *spc15:cas10^HD^* | This study |
| pAS55 | pCRISPR *S. epidermidis* type III-A *spc47:cas10^HD^* | This study |
| pAS61 | mCherry expression. | This study |
| pAS66 | pCRISPR *S. epidermidis* type III-A *spc17.* Used to clone pAS71 | This study |
| pAS67 | pCRISPR *S. epidermidis* type III-A *spc19.* Used to clone pAS72 | This study |
| pAS69 | pCRISPR *S. epidermidis* type III-A *spc29.* Used to clone pAS72 | This study |
| pAS70 | pCRISPR *S. epidermidis* type III-A *spc36.* Used to clone pAS74, pAS78 | This study |
| pAS71 | pCRISPR *S. epidermidis* type III-A *spc17:dcsm6* | This study |
| pAS72 | pCRISPR *S. epidermidis* type III-A *spc19:dcsm6* | This study |
| pAS73 | pCRISPR *S. epidermidis* type III-A *spc29:dcsm6* | This study |
| pAS74 | pCRISPR *S. epidermidis* type III-A *spc36:dcsm6* | This study |
| pAS78 | pCRISPR *S. epidermidis* type III-A *spc36:cas10^HD^* | This study |
| pLM554 | pCRISPR *S. epidermidis* type III-A ∆*spc* | (4) |
| pGG79-F | Used to clone pAS20 | (5) |
| pDB114 | Used to clone pAS22 | (6) |
| pC194 | Used to clone pAS23 | (7) |
| pGG-BsaI-R | Used to clone pAS70 | (8) |
| pFNMB0-mCherry-2 | Used to clone pAS61 | (9) |
| pNA1 | pCRISPR *S. epidermidis* type III-A ∆array, used for spacer library assay | (10) |
| pNA23 | pCRISPR *S. epidermidis* type III-A d*csm6* ∆array, used for spacer library assay | This study |
| pNA24 | pCRISPR *S. epidermidis* type III-A *cas10*^HD^ ∆array, used for spacer library assay | This study |
| pLZ12spec | Used to clone pNA15 | (11) |
| pNA15 | The CRISPR array of *S. epidermidis* type III-A, with a BsaI spacer used for Golden Gate assembly of the spacer library | This study |
| pNA2 | pCRISPR *S. epidermidis* type III-A with single repeat, used for naïve spacer acquisition assay | (10) |
| pNA3 | pCRISPR *S. epidermidis* type III-A d*csm6* with single repeat, used for naïve spacer acquisition assay | This study |
| pNA4 | pCRISPR *S. epidermidis* type III-A *cas10*^HD^ with single repeat, used for naïve spacer acquisition assay | This study |
| pNL03 | pCRISPR *S. epidermidis* type III-A *spc35* | This study |
| pNL09 | pCRISPR *S. epidermidis* type III-A *spc35:dcsm6* | This study |

**Table S6. Plasmids used in this study.**

**REFERENCES**

1. Kreiswirth BN, Lofdahl S, Betley MJ, O'Reilly M, Schlievert PM, Bergdoll MS and Novick RP. (1983) The toxic shock syndrome exotoxin structural gene is not detectably transmitted by a prophage. *Nature*, 305, 709-712.
2. Pyenson NC, Gayvert K, Varble A, Elemento O, Marraffini LA. (2017) Broad Targeting Specificity during Bacterial Type III CRISPR-Cas Immunity Constrains Viral Escape. *Cell Host Microbe*, 22(3), 343-353.
3. Charpentier E, Anton AI, Barry P, Alfonso B, Fang Y, Novick RP. (2004) Novel cassette-based shuttle vector system for gram-positive bacteria. *Appl Environ Microbiol.* 70(10), 6076-85.
4. Hatoum-Aslan A, Samai P, Maniv I, Jiang W, and Marraffini, LA. (2013) A ruler protein in a complex for a ntiviral defense determines the length of small interfering CRISPR RNAs. J*.Biol.Chem.* 288, 27888–27897.
5. Goldberg GW, Jiang W, Bikard D, and Marraffini LA. (2014) Conditional tolerance of temperate phages via transcription-dependent CRISPR-Cas targeting. *Nature*, 514, 633–637.
6. Bikard D, Euler CW, Jiang W, Nussenzweig PM, Goldberg GW, Duportet X, Fischetti VA, and Marraffini LA. (2014) Exploiting CRISPR-Cas nucleases to produce sequence-specific antimicrobials. *Nat. Biotechnol*. 32, 1146–1150.
7. Horinouchi S, and Weisblum B. (1982a) Nucleotide sequence and functional map of pC194, a plasmid that specifies inducible chloramphenicol resistance. *J.Bacteriol.*150, 815–825.
8. Samai P, Pyenson N, Jiang W, Goldberg GW, Hatoum-Aslan A, Marraffini LA. (2015) Co-transcriptional DNA and RNA Cleavage during Type III CRISPR-Cas Immunity. *Cell*, 161(5), 1164-1174.
9. Brodmann M, Heilig R, Broz P, Basler M. (2018) Mobilizable plasmids for tunable gene expression in *Francisella novicida*. *Front. Cell. Infect. Microbiol.* (8)
10. Aviram N, Thornal AN, Zeevi David, Marraffini LA+ (2022). Different modes of spacer acquisition by the Staphylococcus epidermidis type III-A CRISPR-Cas system. Nucleic Acid Research 50, 1661-1672.
11. Husmann, L.K., Scott, J.R., Lindahl, G. and Stenberg, L. (1995) Expression of the Arp protein, a member of the M protein family, is not sufficient to inhibit phagocytosis of Streptococcus pyogenes. *Infect. Immun.,* 63, 345-348.
