## Supplementary material for "The Cas10 nuclease activity relieves host dormancy to facilitate spacer acquisition and retention during type III-A CRISPR immunity": Table S7

| **Name** | **Sequence (5’-3’)** |
| --- | --- |
| AS25 | GATATAAATGGTTTAGCAAATTCCATAGCCGCTAATTTAGATAC |
| AS26 | GTATCTAAATTAGCGGCTATGGAATTTGCTAAACCATTTATATC |
| AS83 | GAACGGAATTTGTATTCTTCTTGAAGTTTTCCTATTAAT |
| AS84 | CGATATTAATAGGAAAACTTCAAGAAGAATACAAATTCC |
| AS94 | AAACTATAATGTTGTTATTTGCTAATAGTTTGTTG |
| AS95 | AAAACAACAAACTATTAGCAAATAACAACATTATA |
| AS96 | GGAGGAATTACAAATGAACGCAC |
| AS97 | GATTCCATTCTTTTGTTTGTCTGCC |
| AS98 | CTGATCAAATACTCTTCCAATTTAG |
| AS99 | GTGCTGAAGTCAAGTTTGAAGG |
| AS100 | GGATAAAGTGGGATATTTTTAAAATCAACCCTCCTCATCACAATGGCAGTTGTGACGTGG |
| AS101 | CCACGTCACAACTGCCATTGTGATGAGGAGGGTTGATTTTAAAAATATCCCACTTTATCC |
| AS102 | CTTTACTCATTGATTAACTCCTCCTTAAAATTATTTGCTAATAGTTTGTTCTGCGAACTT |
| AS103 | TCGCCTTTTGAAGTTCGCAGAACAAACTATTAGCAAATAATTTTAAGGAGGAGTTAATCA |
| AS104 | GCAAATAATTTTAAGGAGGAGTTAATCAATGAGTAAAGGAGAAGAACTTTTCACTGGAGT |
| AS105 | TTGGGACAACTCCAGTGAAAAGTTCTTCTCCTTTACTCATTGATTAACTCCTCCTTAAAA |
| AS106 | CACATGGCATGGATGAACTATACAAATAACAACATTATACACGAAAGGAAAGATAGAAAT |
| AS107 | ATTTCTATCTTTCCTTTCGTGTATAATGTTGTTATTTGTATAGTTCATCCATGCCATGTG |
| AS108 | CCGCAAAGATAATTTAGGTGTAGAAAATTTATACATTGATATATTTATGTTACAGTAATA |
| AS109 | TATTACTGTAACATAAATATATCAATGTATAAATTTTCTACACCTAAATTATCTTTGCGG |
| AS171 | GTATGGCTCTTTATTAGCAGCGATAGGGAAAATTATATATCGAAGTGGTGATCATAC |
| AS172 | GTATGATCACCACTTCGATATATAATTTTCCCTATCGCTGCTAATAAAGAGCCATAC |
| JTR496 | GGCTCTTTATTAGCAGCGATCGGGAAAATTATATATCGAAGTGG |
| JTR497 | CCCGATCGCTGCTAATAAAGAGCCATACATTAATATATTTTTTTTATTCATTTTAACC |
| NA101 | NNNNNAGAAAAGGGGACGAGAAC |
| NA102 | NNNNNTCGGGGTGGGTATCGATC |
| NA134 | CACATCAGAAAATGGAATATCAGGTAGTAATTCC |
| NA135 | GGAATTACTACCTGATATTCCATTTTCTGATGTG |
| NA146 | CTGAAGGAAGATCTGGATCCTAATGAACGGATCCTTTAAAGTATATATCAGATTGTTTCG |
| NA147 | ACTGATGGGCCCCTGCAGATGAATTTTTTCCATCCCCTAGAAATTAATCAATGCGTATTT |
| NA148 | TGATTAATTTCTAGGGGATGGAAAAAATTCATCTGCAGGGGCCCATCAGTCTGACGACCA |
| NA149 | TGATATATACTTTAAAGGATCCGTTCATTAGGATCCAGATCTTCCTTCAGGTTATGACCA |
| NA162 | NNNNNGGCATTTGTTAAAGTATCGGATC |
| NA163 | NNNNNTAGAGTGTTGTTAGATTTAGTTC |
| NA169 | NNNNNTTTGAATGGTCCCA |
| NA170 | NNNNNACAGCATAGCTCTA |
| NA567 | GAACTCGTTAAATAGTCATCATCATTTCTAAAATTGTCCA |
| NA568 | GATCTGGACAATTTTAGAAATGATGATGACTATTTAACGA |
| PS247 | GATATAAATGGTTTAGCAAATTCCATAGCCGCTAATTTAGATAC |
| PS248 | GTATCTAAATTAGCGGCTATGGAATTTGCTAAACCATTTATATC |
| W614 | GGTTATACTAAAAGTCGTTTGTTGG |
| W852 | CCAACAAACGACTTTTAGTATAACC |
| NL41 | GAACAAATAAACTCCACCTGAAATAGTTTTAATTTTAAC |
| NL42 | CGATGTTAAAATTAAAACTATTTCAGGTGGAGTTTATTT |

**Table S7. Oligonucleotides used in this study.**
