## Supplementary material for "The Cas10 nuclease activity relieves host dormancy to facilitate spacer acquisition and retention during type III-A CRISPR immunity": Table S8

| **Plasmid name** | **Cloning strategy** |
| --- | --- |
| pAS04 | pNP54 was used as the backbone in a 2-piece Gibson assembly. AS25-AS26 introduced the *dCsm6* mutations. |
| pAS20 | pGG79-F was digested with BsaI. Oligonucleotides AS83-AS84 were annealed and ligated into the digested backbone to introduce *spc15.* |
| pAS21 | pAS20 was used as the backbone in a 2-piece Gibson assembly. AS25-AS26 introduced the *dCsm6* mutations. |
| pAS22 | pDB114 was digested with BsaI. Oligonucleotides AS94-AS95 were annealed and ligated into the digested backbone to introduce a targeting spacer against wildtype ΦNM4γ4 in the creation of the GFP-NM4 phage. |
| pAS23 | pC194, pCN57 and wildtype ΦNM4γ4 were used as backbones in a 4-piece Gibson to create a plasmid with GFP flanked by ΦNM4γ4 homology arms. The oligonucleotides used were AS96 through AS109. |
| pAS53 | pAS20 was used as the backbone in a 2-piece Gibson assembly. AS171-AS172 introduced the *cas10^HD^* mutations. |
| pAS55 | pNP54 was used as the backbone in a 2-piece Gibson assembly. AS171-AS172 introduced the *cas10^HD^* mutations. |
| pAS69 | pGG79-F was digested with BsaI. Oligonucleotides NA567 and NA568 were annealed and ligated into the digested backbone to introduce *spc29.* |
| pAS73 | pAS69 was used as the backbone in a 2-piece Gibson assembly. AS25-AS26 introduced the *dCsm6* mutations. |
| pAS70 | pGG79-F was digested with BsaI. Oligonucleotides NA567 and NA568 were annealed and ligated into the digested backbone to introduce *spc36.* |
| pAS74 | pAS70 was used as the backbone in a 2-piece Gibson assembly. AS25-AS26 introduced the *dCsm6* mutations. |
| pAS78 | pAS70 was used as the backbone in a 2-piece Gibson assembly. AS171-AS172 introduced the *cas10^HD^* mutations. |
| pNA3 | PCR amplification of pNA2 using PS427/W614 and PS248/W852, followed by Gibson assembly of the two PCR fragments. |
| pNA4 | PCR amplification of pNA2 using JTR496/W614 and JTR497/W852, followed by Gibson assembly of the two PCR fragments. |
| pNA15 | PCR amplification of pGG-BsaI-R using NA146/NA147, and of pLZ12spec with NA149/NA134 and NA148/NA135, followed by Gibson assembly of the three PCR fragments. |
| pNA23 | PCR amplification of pNA1 using PS427/W614 and PS248/W852, followed by Gibson assembly of the two PCR fragments. |
| pNA24 | PCR amplification of pNA1 using JTR496/W614 and JTR497/W852, followed by Gibson assembly of the two PCR fragments. |
| pNL03 | pGG79-F was digested with BsaI. Oligonucleotides NL41 and NL42 were annealed and ligated into the digested backbone to introduce *spc35.* |
| pNL09 | pNL03 was used as the backbone in a 2-piece Gibson assembly. AS25-AS26 introduced the *dCsm6* mutations. |

**Table S8. Plasmid cloning strategies used in this study.**
